# Integrating morphology and gene expression of neural cells in unpaired single-cell data using GeoAdvAE

**DOI:** 10.1101/2025.11.19.689368

**Authors:** Jinqiu Turbo Du, Tom Chartrand, Suman Jayadev, Katherine E. Prater, Kevin Z. Lin

## Abstract

**Background:** Cellular morphological transitions are observed across many diseases, yet their functional role remains unclear because few technologies profile form and function in the same cell. Linking single-cell morphology to transcriptomics is difficult: the two modalities share no feature correspondence and are typically measured in different cells.

**Methods:** We present GeoAdvAE, a geometry-aware adversarial autoencoder for diagonal (unpaired) integration of single-cell morphology and single-cell RNA sequencing. GeoAdvAE couples modality-specific variational autoencoders with a Gromov-Wasserstein regularizer and an adversarial discriminator to embed unpaired morphologies and transcriptomes into a shared latent space that preserves both reconstruction fidelity and cross-modal geometry.

**Results:** Using patch-seq neurons with joint morphology-RNA measurements as ground truth, GeoAdvAE attains the best cross-modal cell-type matching accuracy among diagonal integration methods, outperforming optimal-transport, latent-alignment, and adversarial baselines. Applied to 98 CAJAL-quantified microglial morphologies and 31,948 single-cell transcriptomes from the 5xFAD Alzheimer’s disease model, GeoAdvAE recovers a one-dimensional axis that aligns the two modalities. Integrated-gradient attribution highlights transcriptomic shifts (DNA repair in ramified microglia; cell killing in amoeboid microglia), nominates gene markers (*Ms4a6b*; *Ftl1* /*Fth1*), and reveals disease-associated microglia signatures that are decoupled from morphology.

**Conclusions:** GeoAdvAE provides a scalable and interpretable approach to connecting cellular “form” and “function” when joint profiling of morphology and transcriptomics is impractical. Our method is publicly available at https://github.com/turbodu222/GeoAdVAE.

## 1 Introduction

“Form follows function” is a powerful organizing principle in biology: a cell’s shape is often a visible readout of what it is doing. In the brain, for example, neurons in different layers exhibit distinct arborization patterns that enable specialized circuit roles [32,12,3]. As another example, during the progression of Alzheimer’s disease (AD), microglia remodel their processes and shift their morphology as they surveil tissue, respond to damage, and interact with pathology. Yet morphology is not a perfect proxy for cellular function [39,6]. Similar shapes can conceal divergent molecular states, and conversely, functional reprogramming does not always manifest as a clear change in shape. This gap complicates efforts to understand how microglia respond to disease burden and limits our ability to translate visible phenotypes into mechanistic insight [30].

Molecular profiling provides a complementary view. High-throughput measurements of RNA, proteins, and chromatin capture internal programs and environmental responses at single-cell resolution. However, for many questions, such as the nervous system, where cell morphology is intricate and tightly linked to function, simultaneous measurement of detailed morphology and transcriptomes remains rare. As a result, most available datasets are *unpaired* : large imaging collections with reconstructed shapes and, separately, large single-cell RNA-seq atlases. Unlocking the biology requires principled methods that can connect these modalities without joint measurements.

Integrating morphology with gene expression poses challenges distinct from those of conventional multi-omics alignment. Only a small fraction of genes directly influences morphological features, so information is intrinsically imbalanced across modalities [41,23]. Unlike RNA-ATAC or RNA-protein, there is no straightforward feature-to-feature correspondence to anchor alignment. Moreover, morphology must first be converted into quantitative descriptors that respect geometric relationships, and naïve embeddings can distort them. Together, these issues make diagonal (unpaired) integration between morphology and transcriptomics both necessary and non-trivial.

We introduce **GeoAdvAE**, a geometry-aware adversarial autoencoder for diagonal integration of single-cell morphology and gene expression. GeoAdvAE learns a shared latent space from unpaired datasets using four complementary ingredients: modality-specific autoencoders, an adversarial objective with a Gromov-Wasserstein regularizer for alignment, and a biological prior for orientation. Intuitively, the autoencoders compress each modality into a common low-dimensional space; the adversarial objective encourages morphology and gene-expression cells to overlap within that space; the Gromov-Wasserstein regularizer preserves each modality’s internal geometry, so that cells that are similar in shape (or in expression) remain close after integration; and the biological prior anchors the alignment so that broad, known cell categories correspond across the two modalities. The result is a joint representation that mixes cells across modalities while maintaining biologically meaningful neighborhoods. We validate GeoAdvAE using patch-seq neurons, where matched morphology-RNA measurements provide ground truth for cross-modal cell-type matching. We then deploy the framework to unpaired gene expression and morphology datasets of microglia in the 5xFAD mouse model of AD. By probing the fitted model, we highlight gene programs associated with morphological transitions and identify transcriptomic signatures that are decoupled from visible shape changes. Because GeoAdvAE nominates specific genes and pathways, these computationally derived hypotheses can be prioritized for experimental follow-up in cells (e.g., targeted perturbation) to confirm that they reflect reliable biology.

### 1.1 Biological relevance

Alzheimer’s disease (AD) lacks a cure [47], motivating strategies that enhance resilience – the capacity of cells and circuits to preserve function despite pathology [8]. Microglia are central to this idea: they adopt distinct morphologies that can encircle amyloid plaques and form barrier-like structures associated with tissue protection, even as dystrophic microglia accumulate with aging and may signal impaired responses [39,17]. Much of this knowledge comes from microscopy, but direct links between these visible phenotypes and underlying molecular programs remain limited. To close that gap, we apply GeoAdvAE to the 5xFAD mouse, an amyloid-only transgenic model with reduced between-sample variability. This setting lets us cleanly interrogate RNA-morphology coupling: even if mouse models do not fully reproduce human resilience or dystrophic signatures, they robustly recapitulate a homeostatic-to-inflammatory microglial continuum (historically framed as M2-M1) [16]. By aligning morphology with transcriptomic state in this controlled context, GeoAdvAE provides a roadmap for dissecting putative resilience mechanisms and prioritizing hypotheses for testing in human tissue.

### 1.2 Related work and challenges in integrating morphology and gene expression

Our work addresses the key obstacle that integrating cellular morphology with gene expression is substantially different and arguably more difficult than conventional single-cell integration of two different types, such as RNA-ATAC or RNA-protein integration. We highlight two main reasons for this.

#### Imbalanced information and no clear correspondence

Only a small subset of genes directly influences cell shape (e.g., cytoskeleton, membrane dynamics), so morphology-GEX integration is intrinsically asymmetric and has a low signal-to-noise ratio. This is exemplified in Figure 1, which illustrates patch-seq neurons whose GEX and morphology are measured simultaneously. Neurons with very similar GEX could have vastly different morphologies, and vice versa. By contrast, GEX-ATAC and GEX-protein have natural anchors (i.e., regulatory links and protein readouts) supporting feature-level correspondence [5]. Methods built for such settings (e.g., SCOT [9], cross-modal autoencoders [46]) tend to assume symmetric information and therefore underperform on morphology-GEX. Morphology emerges from many pathways rather than single genes, precluding one-to-one anchors and weakening co-occurrence/paired-feature strategies [5]. We note that quantifying morphology itself is also nontrivial. Tools like CAJAL [13] and MorphOMICs [6] yield embeddings, but links to transcriptional programs remain limited.

**Fig. 1.**
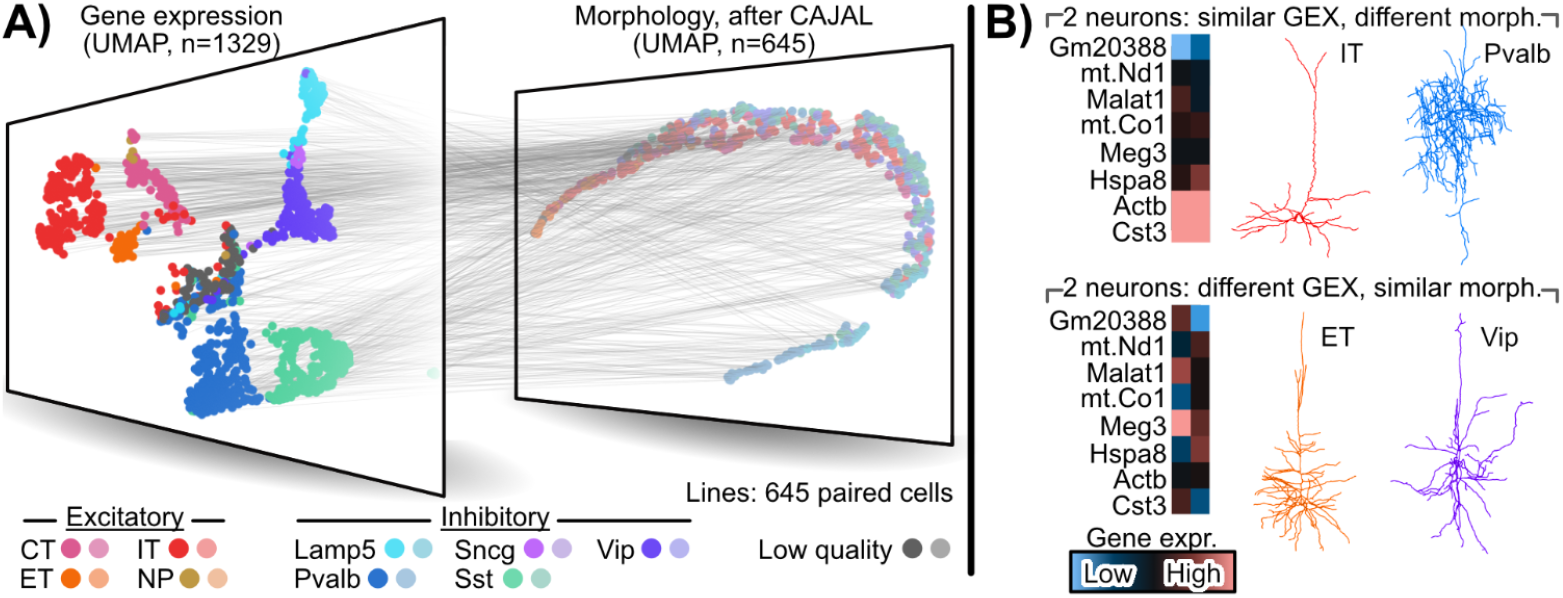
Patch-seq data of cell GEX and morphology. A) Patch-seq data of 1329 neurons [32], of which 645 cells are measured in both gene expression (GEX) and morphology (quantified by CAJAL [13]) among different sub-types of excitatory and inhibitory neurons. The gray lines reveal the complex relation between the two modalities. B) Exemplary genes and morphologies to demonstrate that despite neurons having similar gene expression, they could have vastly different morphologies (top) or vice-versa (bottom).

#### Lack of datasets that measure both genes and morphology of neural (brain) cells

Despite progress in multi-modal integration, linking single-cell morphology with gene expression remains difficult because paired datasets are rare. We note that there is a plethora of work that studies the relationship between gene expression and morphology using spatial transcriptomics [4,19]. However, these spatial transcriptomics platforms lack the resolution to capture the intricate morphology of neural cells [29]. For example, while the latest versions of 10x Visium HD can capture spots of up to 2 *µ*m, this resolution is still not apt for capturing the intricate morphologies of microglia processes that are typically only 1 *µ*m thick [43]. Patch-seq [12] is one of the few wet-bench protocols that enable accurate profiling of both gene expression and brain cell morphology, but its low throughput contrasts with the scale of scRNA-seq and microscopy, which are typically generated separately. Consequently, new frameworks for effective inference from large, unpaired single-cell gene expression and morphology datasets are needed.

## 2 Method

To integrate unpaired cellular morphology and gene expression data, we propose **GeoAdvAE** (geometry-aware adversarial autoencoder), a diagonal integration framework that learns joint representations of both modalities in a shared latent space (Fig. 2A). GeoAdvAE consists of two modality-specific Variational Autoencoders (VAEs) and one discriminator: the two encoders extract features from morphology and gene expression, respectively, and project them into a common latent distribution. Unlike conventional VAEs that rely solely on reconstruction and KL-divergence losses, GeoAdvAE introduces three additional and complementary penalty terms (adversarial alignment, Gromov-Wasserstein-based structural regularization, and a coarse biological cluster-level prior) that cooperatively guide the training process toward desirable cross-modal alignment. Together, these components enable the model to preserve modality-specific fidelity while enforcing semantic and geometric consistency across unpaired single-cell modalities, forming the foundation for integrating morphology and transcriptomic information in the subsequent stages of our framework.

**Fig. 2.**
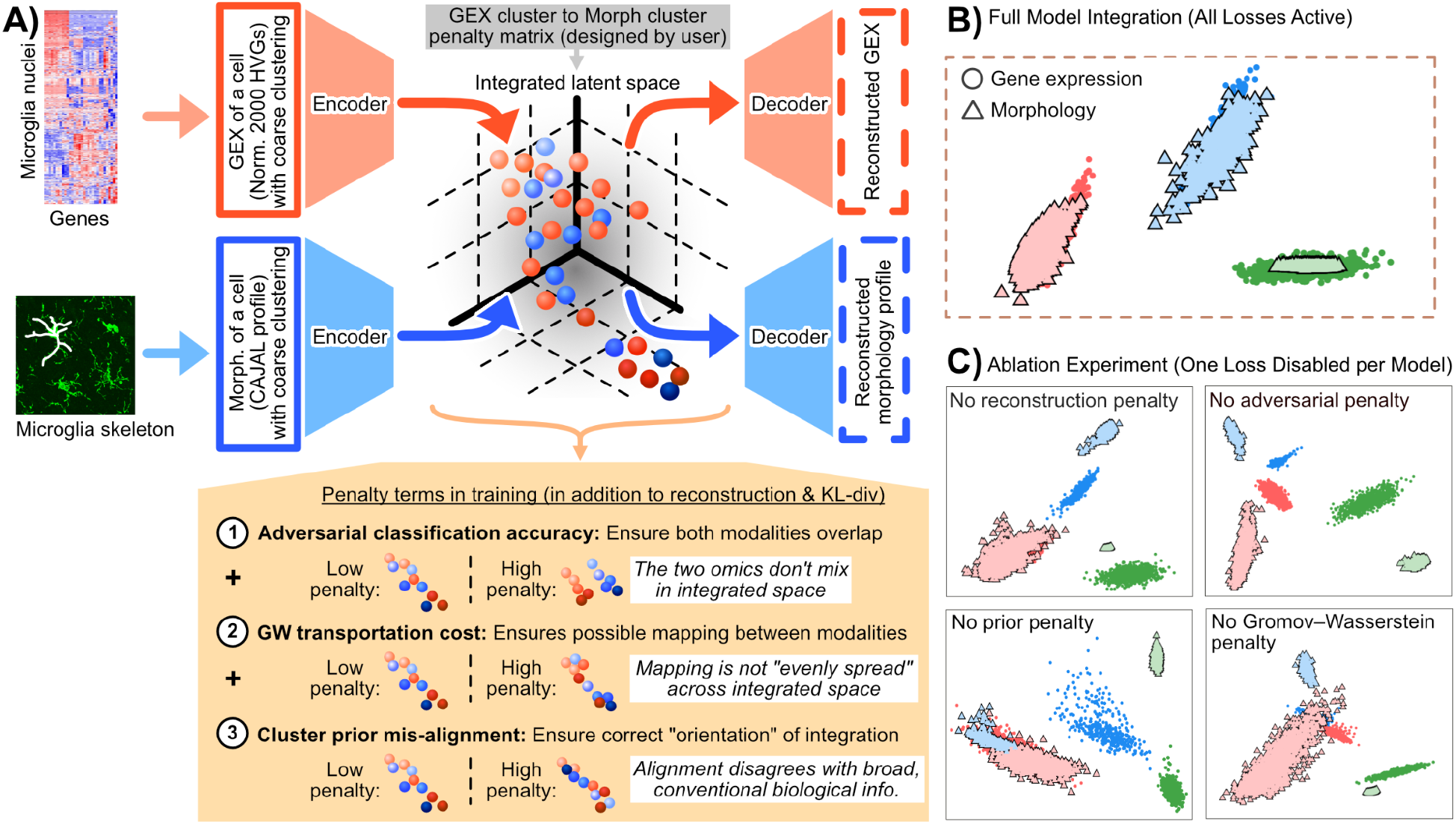
Architecture and ablation study. A) Overview of the GeoAdvAE architecture integrating unpaired morphology (microglial skeleton; white) and gene expression through two modality-specific encoders and decoders in a shared latent space. Three additional loss terms (adversarial alignment, Gromov-Wasserstein (GW) transport regularization, and cluster-prior guidance) jointly control modality overlap, geometric consistency, and biologically meaningful orientation, respectively. The white boxes denote qualitative statements on how to interpret potential integrations with high penalty terms (i.e., an undesirable integration). B) Integrated latent space from simulated unpaired data detailed in Section 4.1 computed by GeoAdvAE. C) Ablation experiment (one penalty disabled for each fit), where removing any of the penalty terms leads to distinct distortions in the integrated latent space. See Appendix S4 for a full Shapley value analysis.

For GeoAdvAE to integrate RNA and morphology, we first need to meaningfully represent a cell’s morphology as a vector of numerical values. To do this, we use CAJAL [13], which represents each microglia’s morphology as a low-dimensional vector based on the Wasserstein distance to other cells’ morphologies. Compared with other methods for quantifying morphology, we have empirically found that CAJAL is the most suitable for our proposed diagonal integration framework (see Fig. S12).

### 2.1 Model architecture

#### Encoder and decoder architecture

Each modality uses its own VAE. We denote gene expression as modality *A* and morphology as modality *B*, with modality-specific encoders *E*_*A*_ and *E*_*B*_, respectively. The gene expression profiles (**x**_*A*_ ∈ ℝ^2000^, for the 2000 highly variable genes, log-normalized) are organized into three layers. Both encoders output the mean and log-variance of a *d* = 16 integrated latent space. The morphology profiles (**x**_*B*_ ∈ ℝ^30^, quantified via CAJAL [13] as described in Section 3) pass through two hidden layers. Using the reparameterization trick, let **z**^(*A*)^ and **z**^(*B*)^ be sampled from the posterior Gaussian distributions produced by the gene expression encoder *E*_*A*_ and morphology encoder *E*_*B*_ using **x**_*A*_ and **x**_*B*_ as inputs, respectively. Decoders mirror the encoders back to the input size with the same blocks; the final layer is linear to preserve real-valued reconstructions.

#### Discriminator architecture

To align the latent spaces across modalities, we add a discriminator *D* : ℝ^*d*^ → [0, 1], which is a three-layer multi-layer perceptron. The encoders are updated adversarially to fool *D*, yielding a shared, modality-invariant latent.

### 2.2 Loss Function Formulation for Cross-Modal Alignment

We align gene expression (GEX, modality *A*) and CAJAL morphology embeddings (modality *B*) in a shared latent space through a weighted sum of five complementary objectives:

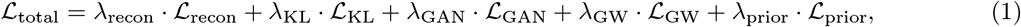

with hyperparameters *λ*_KL_, *λ*_recon_, *λ*_prior_, *λ*_GAN_, *λ*_GW_ ≥ 0. Our default choices are *λ*_KL_ = 0.1, *λ*_recon_ = 5, *λ*_prior_ = 1, *λ*_GAN_ = 4, and *λ*_GW_ = 1. Training follows a curriculum: autoencoding (*L*_KL_+*L*_recon_) is active from the start; the adversarial term (*L*_GAN_) is enabled after an initial warmup so that encoders learn stable reconstructions first; the prior-guided semantic alignment (*L*_prior_) ramps up via a schedule; finally, the GW structure term (*L*_GW_) is enabled after the latent geometry stabilizes. As our results in Section 4 will emphasize, integration that focuses solely on reconstruction is insufficient to integrate GEX and morphology. The addition of these other penalty terms encourages GeoAdvAE to prioritize other qualities that enable meaningful biological discovery beyond what existing methods optimize for.

#### Reconstruction loss and KL divergence

For each modality, the encoder parameterizes a diagonal-Gaussian posterior; we apply the Kullback-Leibler (KL) divergence penalty against a unit-normal prior to regularize the encoder. Each modality’s decoder reconstructs its original input; we use an *L*_1_ loss (Manhattan distance) to measure the reconstruction quality for robustness in high dimensions. Both the reconstruction and KL divergence penalties are summed across both modalities.

#### Adversarial classification based on the discriminator to ensure integration

Leveraging our discriminator, we include an adversarial classification term to encourage the embeddings from both modalities to overlap. This ensures mixing between the two modalities. Specifically, let *q*_*A*_ and *q*_*B*_ denote the posterior distributions produced by *E*_*A*_ and *E*_*B*_, respectively, and **z**^(*A*)^ and **z**^(*B*)^ be samples from these distributions. The discriminator *D* : ℝ^*d*^→ [0, 1] tries to identify the modality based on the samples **z**^(*A*)^ and **z**^(*B*)^, while encoders try to produce posterior distributions that fool the discriminator.

To train, we alternate between training the discriminator and the generator. The discriminator minimizes the following classification loss with a fixed hyperparameter *ϵ* = 0.1 for stability, where a prediction closer to 0 or 1 means the discriminator predicts the cell to originate from modality *A* or *B*, respectively.

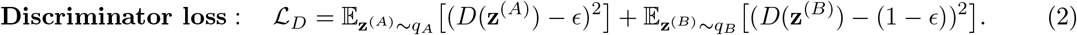

Training the discriminator *D* (but keeping the encoders fixed, and hence, the embeddings **z**^(*A*)^ and **z**^(*B*)^ fixed) to obtain a near-0 would reflect a clear separation between the cell morphology and GEX embeddings, which is not desirable for our integration goal (hence, adversarial).

The generator (i.e., encoders) has a penalty term to encourage better integration between GEX and morphology by updating the encoders (but holding the discriminator fixed),

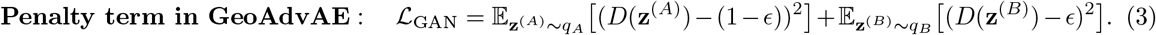

This particular formulation is inspired by Crossmodal-AE [46], where the updated encoder (and hence, updated embeddings **z**^(*A*)^ and **z**^(*B*)^) are incentivized to cause the discriminator to mispredict whether the cell’s embedding originates from the GEX or the morphology modality.

#### Gromov-Wasserstein loss to enable uniform alignment

We align intra-modality geometry by minimizing the Gromov-Wasserstein (GW) discrepancy between pairwise distances in the two latent spaces. This penalty term encourages the integration to spread cells uniformly across the integration space so that as much of a one-to-one mapping between cells from each modality as possible can be achieved. To define this, we first compute the intra-modality distances,

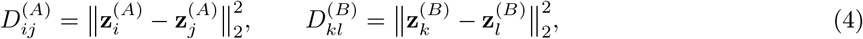

for all cell pairs {*i, j*} and {*k, l*} in modality *A* or *B*, respectively. Then, we match the geometry (among the cells in the minibatch) through an optimal transport plan *T*, a coupling matrix whose entry *T*_*ik*_ specifies the transported mass between cell *i* in modality *A* and cell *k* in modality *B*. We optimize over all valid transport plans *T* ∈ *Π*, where *Π* is the set of doubly stochastic matrices (i.e., rows and columns sum to 1).

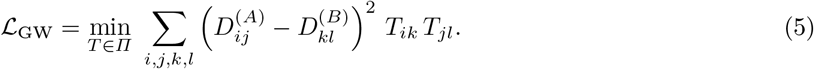

#### Prior-guided cluster alignment

To orient the shared latent space with coarse biology (e.g., excitatory vs. inhibitory in patch-seq), we impose a prior on broad cluster-cluster correspondences. Let *C*_*A*_ : {1, …, *N*_*A*_} → {1, …, *K*_*A*_} and *C*_*B*_ : {1, …, *N*_*B*_} → {1, …, *K*_*B*_} be precomputed broad cluster labels for gene expression (modality *A*) and morphology (modality *B*), where *N*_*A*_ and *N*_*B*_ are the number of cells and *K*_*A*_ and *K*_*B*_ the number of clusters in each modality, respectively. For instance, in our patch-seq analysis, our clustering is simply separating excitatory from inhibitory neurons. In this example, there are well-established distinctions between these two categories of neurons from both gene expression and morphological data. The clusterings we use here are *not* about more granular types of neurons, since they are more difficult to define from a single modality alone. It ensures that broad, literature-supported cell categories remain coherently aligned across modalities, providing semantic structure that complements the unsupervised adversarial and geometric objectives.

The user also provides a correspondence matrix 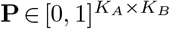 whose entry *P*_*jk*_ encodes the expected association strength between GEX cluster *j* and morphology cluster *k*. This induces a target similarity

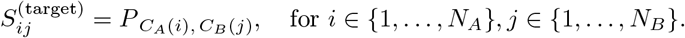

For a minibatch with latent codes 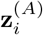 and 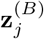, we define the predicted cross-modal similarity by cosine similarity of *ℓ*_2_-normalized embeddings,

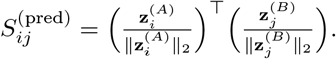

The prior loss matches these similarities via a mean-squared error scaled by a temperature *τ >* 0 that sharpens contrasts, emphasizing confident matches and down-weighting weak ones,

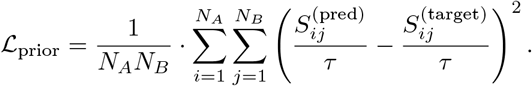

## 3 Data processing and evaluation

The model operates on two unpaired modalities: morphology and gene expression. The morphologies are “skeletons” of the cell in `.swc` format, recording the 3-dimensional location and connection between each “point.” Concretely, a skeleton traces the centerline (rather than the outline) of the cell’s soma and processes as a connected tree of sampled points; each point stores its 3D coordinate and a local radius and links back to a single parent point, and consecutive points are joined by short straight segments that together approximate the smoothly curving processes. These morphologies are processed using CAJAL [13], which encodes structural information in a continuous, numerical space suitable for GeoAdvAE’s deep-learning framework. The transcriptomics are highly variable genes (HVGs, default: 2000), which we log-normalize to remove the confounding effect of sequencing depth.

### Diagonal integration methods to compare against

We benchmark GeoAdvAE against representative cross-modal single-cell integrators spanning distinct paradigms. For instance, coupled/graph autoencoders or latent embedding methods learn shared spaces for unmatched modalities (ScDART [50]; STACI [49]; scJoint [26]). Some methods also incorporate adversarial terms to encourage a better integration (CycleGAN [51], sciCAN [45], SCIM [35], Crossmodal-AE [46]). We emphasize that several of these baselines rely on structure that is absent in our setting: scJoint, sciCAN, and ScDART were built for unpaired scRNA-seq and scATAC-seq integration, where a gene-activity bridge places both modalities on a shared gene axis, and STACI was developed for spatially co-registered transcriptomics and imaging. Because morphology and gene expression share no such gene-activity bridge or spatial graph, we present these four methods as “-like” baselines rather than literal applications of the originals (see Appendix S3.5); the remaining baselines require no feature-correspondence or spatial prior and are applied directly. Regardless, existing diagonal integration methods apt for gene-morphology integration rely on an implicit assumption that the axes of variation that best represent the modality are also the axes shared between the two modalities. As we have seen in patch-seq data (Fig. 1A), this assumption is not necessarily true for GEX-morphology alignment. In contrast, optimal-transport (i.e., geometry matching) methods align the global structures between the two modalities (SCOT [9], MMD-MA [27], UnionCom [2]). However, these methods implicitly assume that the global structure for each modality has similar structural properties, and they also lack the ability to distinguish the “proper orientation” of the integration. The patch-seq data (Fig. 1A) demonstrates that both drawbacks are detrimental when aligning GEX-morphology of neural cells.

### Evaluation criteria

We evaluate diagonal integration via cross-modal cell-type transfer accuracy. These “true” cell-type labels are *not* used during the training of any of the proposed methods. For each query cell in one modality, we take its 1-nearest neighbor (in Manhattan distance) in the other modality within the latent integrated space and count a match if cell-type labels agree. Accuracy is defined as

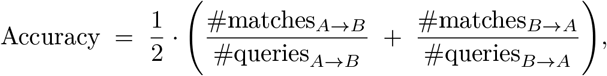

where higher values indicate a better cross-modal alignment.

### Downstream gene investigation

The importance of GeoAdvAE’s diagonal integration lies in its ability to refine our biological understanding of which genes are associated with morphological changes. We use integrated gradients [38], a computational framework for assessing how perturbing a gene’s expression alters the embedding produced by the GEX encoder *E*_*A*_(·). By interpreting each gene’s impact as its “importance scores,” we can identify pathways that concurrently have high importance scores via a gene set enrichment analysis (GSEA, [37]). This downstream analysis can also identify pathways known to be differentially expressed across cell populations that do *not* have a morphological component.

## 4 Results

### 4.1 Simulated data demonstrates GeoAdvAE’s advantage over other methods and enables ablation studies

We built a simulator that generates synthetic gene-expression profiles and corresponding neuronal morphologies to evaluate GeoAdvAE under controlled conditions. This enables us to precisely diagnose how each loss component of GeoAdvAE contributes to its superior performance over other integration methods. Three canonical neuron types (pyramidal, multipolar, bipolar) are modeled by sampling GEX from three low-dimensional clusters; morphology is then generated by a process in which GEX controls polarity, branch density, and anisotropy, yielding three separable CAJAL morphology clusters (Supplemental Fig. S1). For diagonal integration, we use a 3*×*3 correspondence matrix **P** (the identity), which maps each GEX cluster to its morphological counterpart (Fig. 3B).

**Fig. 3.**
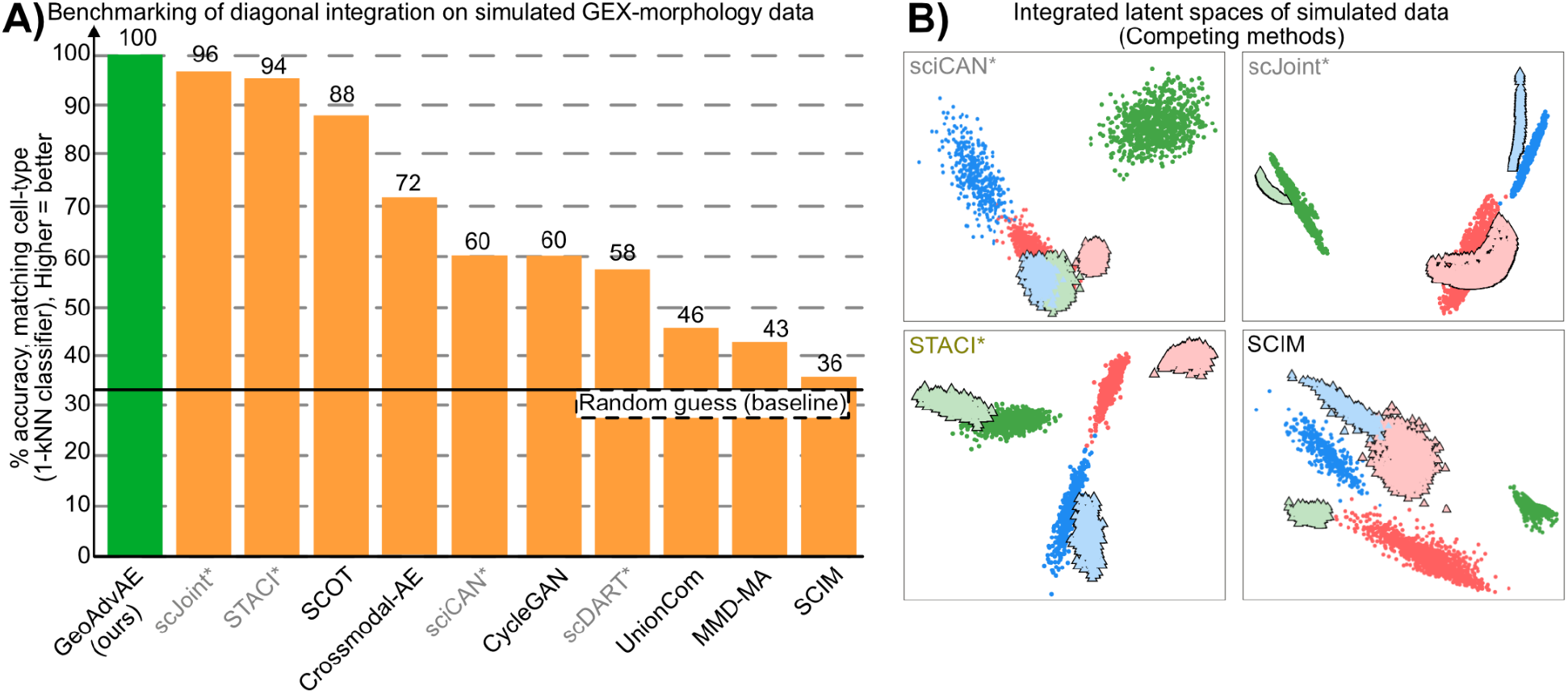
Simulation study via synthetic neurons. A) Comparison of cell-type matching accuracy on simulation data across GeoAdvAE and competing cross-modal integration methods. The solid horizontal line denotes the accuracy achieved by random guessing. B) Integrated latent spaces of simulated data produced by selected baseline methods (sciCAN [45], STACI [49], scJoint [26], SCIM [35]). The results by these competing methods can be compared against GeoAdvAE’s integration shown in Figure 2B. Method names shown in gray and starred denote “-like” baselines: methods that rely on a gene-activity or spatial prior absent from our setting (scJoint, STACI, sciCAN, scDART), as detailed in Section 3 and Appendix S3.

To illustrate the importance of each penalty term in GeoAdvAE, we performed an ablation study, where we removed one of the penalty terms and assessed how the resulting integration suffers (Fig. 2C). Removing the adversarial classifier separates modalities in the latent space. Removing the Gromov-Wasserstein penalty breaks local geometric coherence. Removing prior-guided cluster alignment misorients matched clusters. Removing reconstruction loss destabilizes embeddings and blurs cluster boundaries. Although our correspondence matrix **P** “reveals” how the two modalities should be integrated, we found that this biological information is *necessary* for GEX-morphology integration. In the presence of no paired samples and no matched features between the two modalities, the information represented by **P** provides the necessary orientation signal that the data alone cannot recover.

We next compared GeoAdvAE with other competing methods using both quantitative and qualitative evaluations. GeoAdvAE achieved the highest alignment accuracy (Fig. 3A). Latent embedding methods (scJoint and STACI) performed reasonably well but tended to under-align fine-grained structures, while optimal-transport methods (UnionCom, MMD-MA) struggled to uncover the correct integration orientation (Fig. 3B, Supplemental Fig. S6). As noted in Section 3, scJoint and STACI are adapted “-like” baselines here, since they were originally designed for other modality settings rather than for unpaired morphology-gene expression integration. We also assessed GeoAdvAE in a more complex simulation with five canonical neuron types where the correspondence matrix **P** only offers partial information to separate these types (Supplemental Fig. S2). Collectively, these results support combining adversarial alignment with cluster-prior regularization to achieve proper alignment between GEX and morphology. See Appendix S4 for more results, such as a more in-depth ablation analysis via Shapley values and a comparison between using GW and Maximum Mean Discrepancy (MMD) losses to measure alignment.

### 4.2 Patch-seq neurons provide validation for GeoAdvAE’s integration

As mentioned in Section 1, patch-sequencing is one of the few wet-bench protocols where the GEX and morphology of neurons can be simultaneously measured. This data provides an invaluable resource for validating the performance of any GEX-morphology diagonal integration method. We evaluated our method on a patch-seq dataset [32], focusing on the 645 neurons spanning a diverse set of excitatory and inhibitory neurons from the mouse motor cortex for which both gene expression and morphology were profiled. To define the cluster prior matrix in the patch-seq dataset, we *do not* use the provided cell annotations, as they were defined by the original authors across multiple modalities. Instead, for GEX, we clustered the neurons into 4 clusters and annotated each cluster as either excitatory or inhibitory using canonical marker genes [24]. For morphology, we embedded the neurons into a lower-dimensional space using CAJAL [13] and grouped them into 4 clusters. Based on our visual assessment of 10 randomly selected neurons in each cluster, we annotated whether each neuron was excitatory or inhibitory, and whether it had a pyramidal shape. Based on these manual annotations, we constructed the correspondence matrix **P** mapping between the GEX and morphology clusters based on how highly our coarse excitatory-vs.-inhibitory labels align between the two modalities. The details are described in Appendix S3.

We validated GeoAdvAE using patch-seq neurons with joint GEX-morphology measurements. GeoAdvAE achieved 34% cell-type alignment accuracy, outperforming graph-based baselines such as ScDART (28%) and STACI (27%), as shown in Fig. 4A. Graph-based approaches (GeoAdvAE, ScDART, STACI) performed best, whereas latent-space alignment methods (e.g., scJoint, Crossmodal-AE) and adversarial models (e.g., CycleGAN, sciCAN) degraded on patch-seq, likely because they struggle to preserve modality consistency in the presence of biological noise and nonlinear morphological variability. Consistent with our simulations (Fig. 3A), optimal transport methods (SCOT, UnionCom) performed poorly, as the high diversity of GEX axes often lacks corresponding morphological features. UMAP visualizations confirmed that GeoAdvAE produced a smoother, more biologically consistent latent mixture than Crossmodal-AE (Fig. 4B,D). Additionally, the continuous latent space (Fig. 4B) suggested an alignment of cellular states rather than individual cells (analysis and results shown in Fig. S11). We further benchmarked GeoAdvAE against Supervised PCA [1] to quantify the specific impact of prior information (Fig. S13) and against Tilted-CCA [25] to establish the theoretical “ceiling” for shared signal recovery (Fig. S15).

**Fig. 4.**
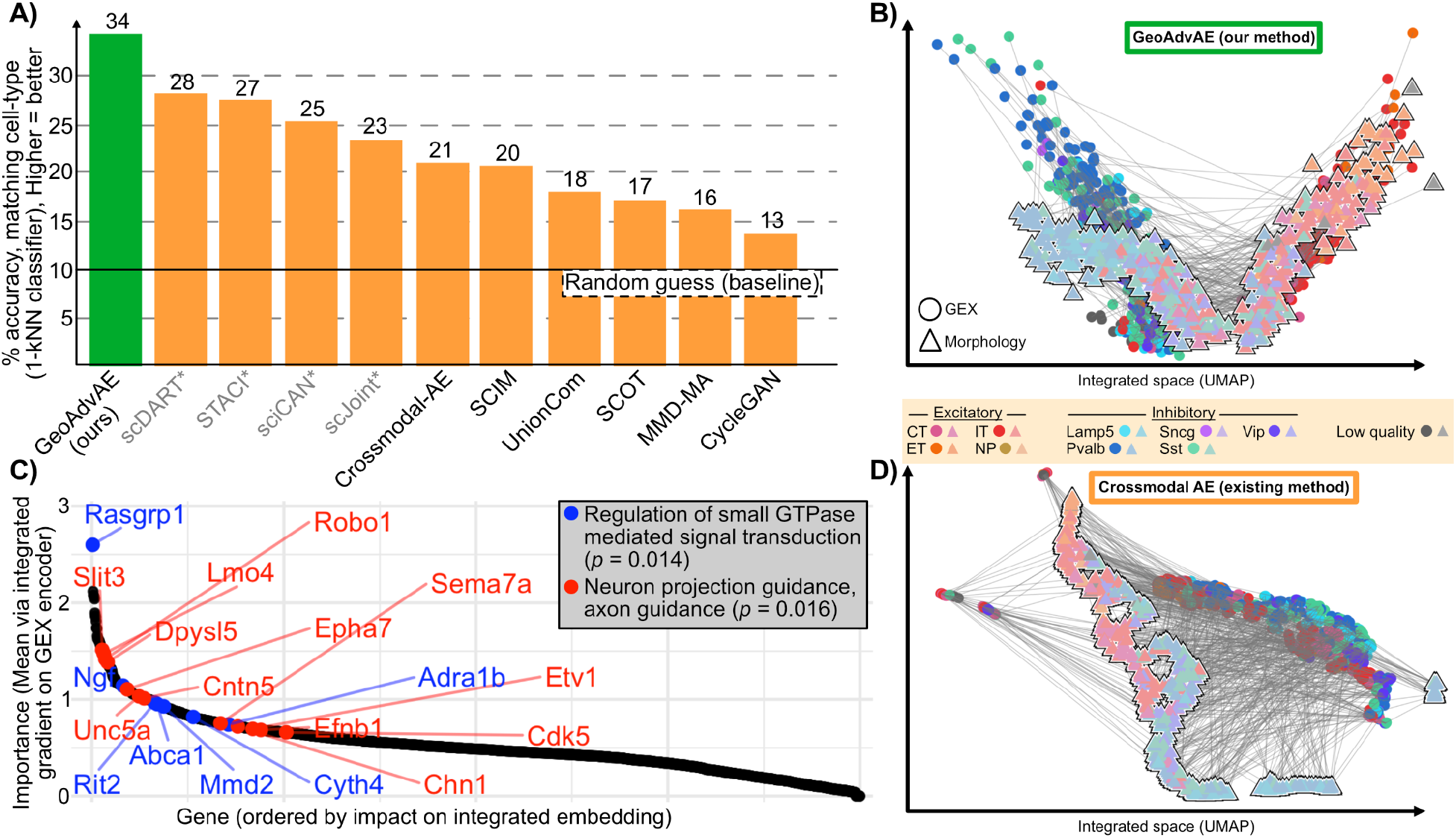
Comparison and validation of integration methods using patch-seq neurons. A) Comparison of cell-type matching accuracy on patch-seq data across GeoAdvAE and competing cross-modal integration methods. GeoAdvAE achieved the highest alignment accuracy. B) Integrated latent space learned by GeoAdvAE on patch-seq neurons. C) Integrated gradient importance scores of all the 2000 genes based on the GEX encoder. The genes in two critical pathways known to regulate neuron morphologies are highlighted in red and blue, both nominally significant. D) Integrated latent space generated by the existing Crossmodal-AE method [46], showing less continuous cross-modal alignment compared to GeoAdvAE. B and D share the same cell-type legend. As in Fig. 3, method names shown in gray and starred denote “-like” baselines that we adapted from settings other than unpaired morphology-gene expression integration (see Section 3 and Appendix S3).

Model interpretation via integrated gradients on the GEX encoder *E*_*A*_(·) confirmed that GeoAdvAE captures essential neuronal biology. We highlight two important pathways identified by GSEA in Fig. 4C.

First, pathways for neuron projection guidance (GO:0097485) and axon guidance play a critical role in axon/neurite guidance and cytoskeletal regulation (example genes: *Slit3* ; p-value: 0.016). Additionally, Rho-family small GTPases are master regulators of neuronal morphogenesis, coupling extracellular cues to actin remodeling that governs spine structure, filopodial dynamics, and growth-cone advance or collapse [28]. In diverse contexts, guidance and trophic signaling converge on these GTPases to control growth-cone behavior and neurite extension [42,7]. Consistent with this framework, we found that the relevant GO term is strongly associated with GEX-morphology, via genes such as *Rasgrp1* and *Ngf*. We ensure these findings are supported by analyses that directly leverage the paired nature of patch-seq data (Fig. S9).

Additional results are in Appendix S4, where we also map the full 1329 GEX cells to the 645 cells with morphology, and also assess how GeoAdvAE performs with classically used morphological features instead of with CAJAL-derived features.

### 4.3 GeoAdvAE uncovers novel biology of 5xFAD microglia

Our primary dataset of interest is one generated on microglia from the 5xFAD mouse model, which recapitulates certain hallmarks of Alzheimer’s disease (AD), see Section 1.1. For morphological data, we leverage 98 microglial skeletons from mice at 3 and 6 months of age and both sexes [6]. After applying CAJAL and clustering, we observe 3 primary microglial clusters (Fig. 5A), where one microglial cluster (cyan) is enriched for larger microglia with a more ramified morphology primarily from male, younger mice. In contrast, another microglial cluster (pink) is enriched for smaller microglia with a more amoeboid morphology, primarily from female, older mice. For transcriptomic data, we leverage 31,948 microglial single-nucleus RNA-sequencing profiles [40]. These microglia span many different states that the authors classified using marker genes, including homeostatic, proliferating, interferon-response (IRM), and disease-associated microglia (DAM) (Fig. 5B).

**Fig. 5.**
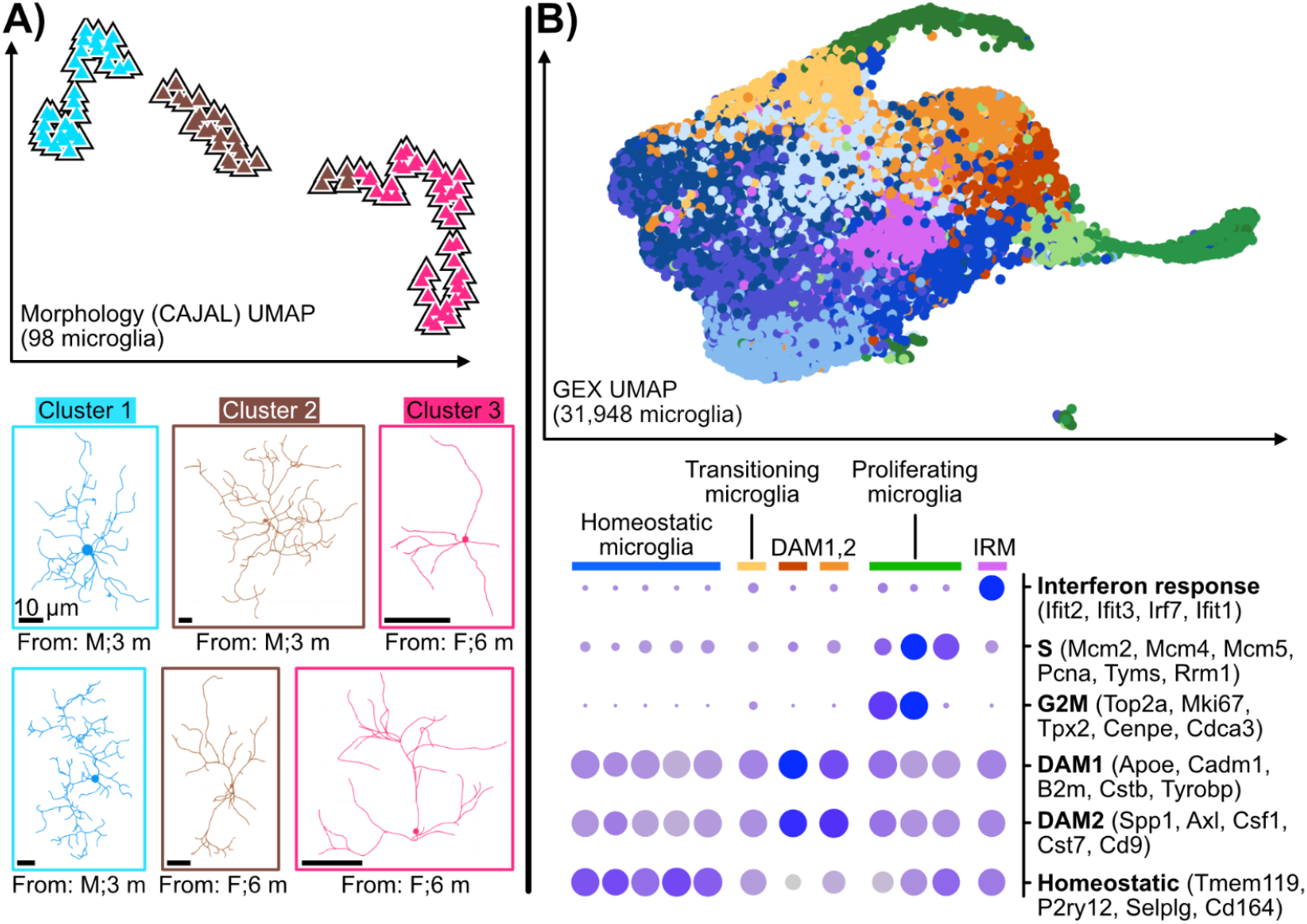
5xFAD microglia data. A) Microglia morphologies [6], quantified via CAJAL and clustered into 3 major morphological types. Exemplary microglia of each cluster are shown below. B) Microglia gene expression [40], using the authors’ annotated microglial states. Marker genes for each microglial state are shown as a dot plot below.

To form the correspondence matrix **P** necessary for the prior-guided cluster alignment, we annotate broad correspondences between the two modalities. For GEX, we use the provided 12 GEX microglia clusters and further quantify the proportions of homeostatic, proliferating, IRM, and DAM microglia within each cluster using marker-gene enrichment analysis. This enables us to capture continuous state transitions among microglial states, rather than discretely labeling each cluster as a “pure” state. For morphology, we clustered the CAJAL profiles into 3 groups and labeled them as ramified, intermediate, and amoeboid based on visual inspection of microglia in each group. Based on these annotations, we design **P** to account for within-cluster proportions and the biological correspondence between microglial transcriptomic and morphological states. See Appendix S3 for more details on this procedure.

When we applied GeoAdvAE to the microglia datasets, we observed an apparent 1-dimensional manifold, suggesting novel biological insights into microglia. Fig. 6A depicts the integrated embedding where the ramified morphologies (cyan triangles) integrate with the homeostatic microglia (blue circles), while the amoeboid morphologies (pink triangles) integrate with the DAM microglia (orange circles). While this overall trend is not surprising due to GeoAdvAE’s prior-guided cluster alignment, we perform downstream analyses to investigate two notable biological questions: 1) which particular genes contributed to this alignment, and 2) what is the continuum of microglial states that spanned this 1-dimensional manifold?

**Fig. 6.**
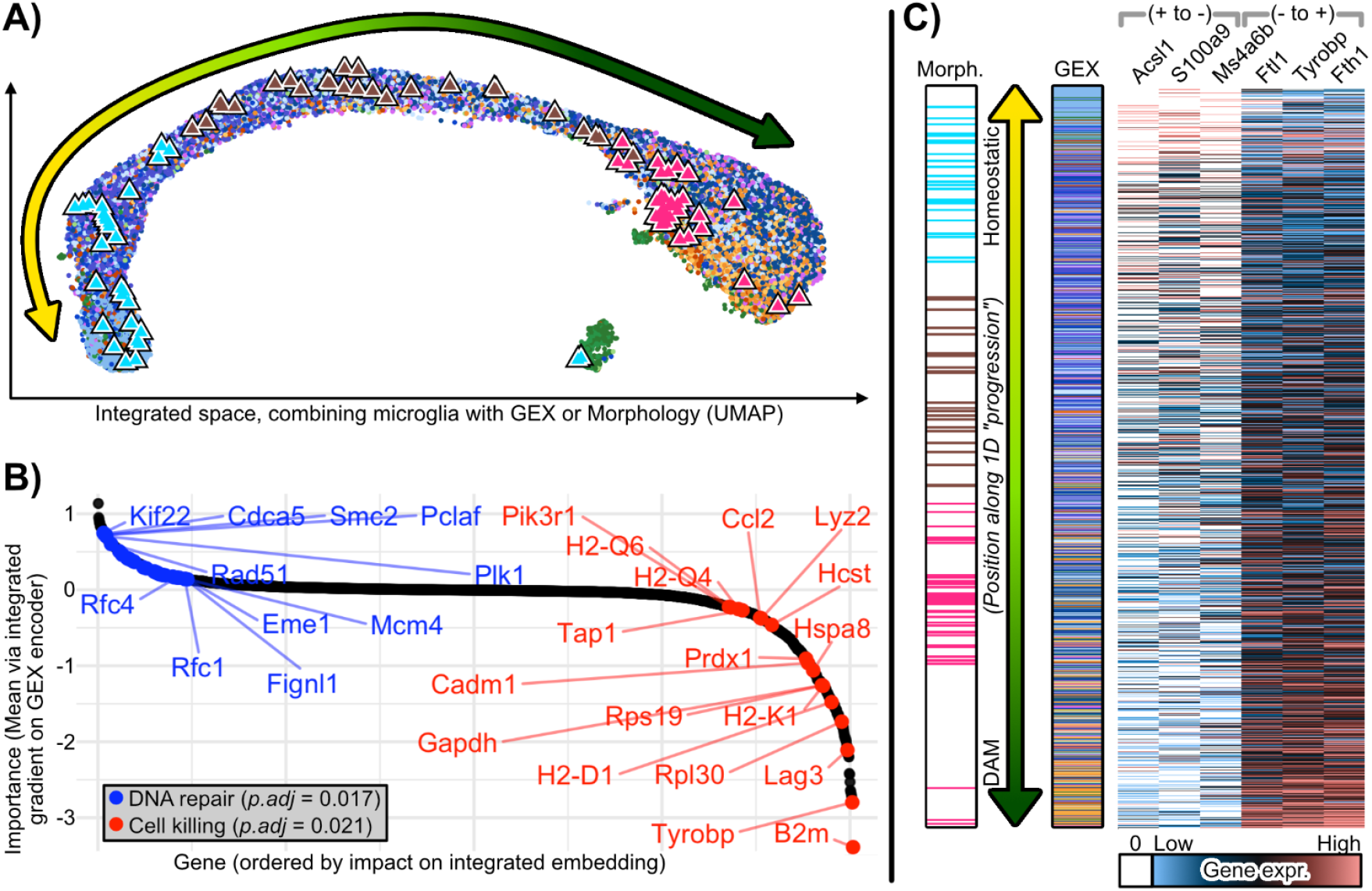
Integration of GEX and morphology uncovers functional roles of microglia in 5xFAD. A) Integration of microglia GEX and morphology, uncovering a 1-dimensional manifold that corresponds to a spectrum of microglial states. B) Genes ordered by their influence on a microglia’s embedding in the integrated space as measured by integrated gradients [38] highlight two major pathways corresponding to the two different ends of the GEX-morphology integrated space. C) Ordering microglia along the 1-dimensional manifold, highlighting how microglia measured by morphology (left) or GEX (right) align along this manifold. Exemplary genes that also generally increase or decrease going from one end of the spectrum to the other are displayed as a heatmap.

We used our integrated gradient framework to identify genes associated with microglial morphology. Ordering genes by importance reveals two significant, opposing pathways (Fig. 6B): DNA repair genes were enriched among ramified microglia, whereas cell-killing genes were enriched among amoeboid microglia. This is consistent with prior work showing that homeostatic, ramified microglia engage in tissue-maintenance programs [30,48,33] while amoeboid microglia adopt highly phagocytic, neurotoxic phenotypes that surround and eliminate stressed or dying neurons [30,10].

To examine genes varying along this continuum, we ordered microglia (both morphology and transcriptomic) along the manifold using Slingshot [36] and correlated each gene with this ordering. We found *Ms4a6b* to be strongly associated with ramified morphologies. This is noteworthy since the closest human homolog, MS4A6A, has conflicting evidence regarding AD. MS4A6A is implicated in disease progression through TREM2 in some studies [31,20] but interpreted as a marker of accelerated aging in others [34,22]. Our results suggested *Ms4a6b* is not primarily linked to inflammatory responses in AD. In contrast to the ramified-associated *Ms4a6b*, we found *Ftl1* and *Fth1* to be associated with amoeboid morphologies, two iron-loading genes whose upregulation is a hallmark of dystrophic microglia, and whose elevated expression localizes around AD plaques and correlates with pathological burden [15,21]. Surprisingly, several disease-associated microglia (DAM) complement markers were not correlated with this GEX-morphology axis, including *C1qa, C1qb, C1qc, C3*, and *C4b* [18,11]. This suggested that complement activation represents an upstream or partially orthogonal transcriptomic program that can be engaged without large shifts in soma size or process complexity, consistent with recent work showing that microglial morphology is only one particular readout of a microglia’s function [14,44]. Future gene knockout experiments could be performed to validate these findings. All in all, these findings demonstrate GeoAdvAE’s ability to uncover promising GEX-morphology relations, enabling us to better characterize the morphological consequences of up/down-regulation of particular microglial transcriptomic programs.

## 5 Discussion

GeoAdvAE provides a general framework for integrating cellular morphology and gene expression from unpaired data, enabling us to link “form” and “function” even when these modalities are measured in different cells. Using simulations and patch-seq neurons, we showed that combining adversarial alignment, Gromov-Wasserstein regularization, and prior-guided cluster correspondences yields more accurate and biologically coherent cross-modal embeddings than existing integration methods. Applied to 5xFAD microglia, GeoAdvAE uncovers a one-dimensional ramified-to-amoeboid continuum, highlights DNA repair and cell-killing programs associated with morphological transitions, and reveals complement/DAM signatures that appear decoupled from visible shape changes. We emphasize that this one-dimensional axis reflects the resolution afforded by only 98 reconstructed microglia rather than a claim that microglial state is intrinsically one-dimensional. Because morphological reconstruction from microscopy remains low-throughput, a single dominant ramified-to-amoeboid axis is the most we can reliably discern at this sample size; richer, higher-dimensional structure that is consistent with the broader heterogeneity now appreciated in microglial biology may well emerge as larger paired morphological datasets become available. These results suggest that some microglial transcriptional programs lie upstream of, or orthogonal to, large-scale morphological remodeling, underscoring the limits of morphology alone as a proxy for microglial state. Because GeoAdvAE requires no explicit cell matching, a natural next step is to evaluate it on single-cell-resolution spatial transcriptomics paired with histological imaging, which could validate the recovered form-function axes in situ and extend the framework to the spatial context of Alzheimer’s disease pathology. More broadly, GeoAdvAE offers a scalable template for integrating unpaired morphology with high-dimensional omics data in other brain and non-brain systems where joint profiling remains impractical.

## Acknowledgments

We thank Juan Caicedo, Elizabeth Haynes, and Sündüz Keleş, the anonymous RECOMB reviewers, and other members of the Lin Lab for support, helpful discussions, and feedback on this work.

## Funding

Research reported in this manuscript was supported by the University of Washington’s Alzheimer’s Disease Research Center (ADRC)’s Development Project and the National Institute of General Medical Sciences (NIGMS) of the National Institutes of Health (NIH) under award number R35GM162089. The content is solely the responsibility of the authors and does not necessarily represent the official views of the National Institutes of Health.

## Author contributions

Jinqiu (Turbo) Du: Conceptualization, Methodology, Software, Formal analysis, Investigation, Visualization, Writing – original draft. Tom Chartrand: Data curation, Methodology (morphology and patch-seq). Suman Jayadev: Methodology (microglia), Writing – review and editing. Katherine E. Prater: Conceptualization (microglial biology), Validation (biological interpretation), Writing – review and editing. Kevin Z. Lin: Conceptualization, Methodology, Supervision, Funding acquisition, Project administration, Writing – original draft, review and editing.

## Conflicts of interest

The authors declare no potential conflicts of interest with respect to the research, authorship, and publication of this article.

## Ethical approval

This study is a secondary computational analysis of previously published, publicly available datasets and did not involve any new experiments on human participants or animals by the authors. Ethical approval was therefore not required for this study. The original patch-seq, 5xFAD microglial morphology, and 5xFAD transcriptomic datasets were generated under the approvals reported in their respective primary publications.

## Consent

Not applicable. This study did not involve new human participants or individual-person data.

## Notes

https://github.com/turbodu222/GeoAdVAE

### Competing Interest Statement

The authors have declared no competing interest.

### Summary of Updates

We have incorporated feedback after thorough discussions from patch-seq and microglia experts, added more extensive simulations, and did a thorough revision of the paper (formatting, typos, clarifications).

