## Supplement for "Integrating morphology and gene expression of neural cells in unpaired single-cell data using GeoAdvAE"

#### S1 Code and data reproducibility

Our code is open source and available at `https://github.com/turbodu222/GeoAdVAE`. This repository includes the implementation of GeoAdvAE, the simulation of GEX-morphology neurons, and the analysis, plots, and benchmarking of our simulated data, patch-seq neurons, and 5xFAD microglia.

The datasets used in this study were obtained from publicly available sources. patch-seq gene expression and morphology data were downloaded from the Berens Lab Mini-Atlas repository (`https://github.com/berenslab/mini-atlas`), originally published by [16]. For the 5xFAD mouse model, the reconstructed neuronal morphologies were obtained from NeuroMorpho (`https://neuromorpho.org/KeywordResult.jsp?keywords=%22siegert%22`), as reported by [4]. The corresponding 5xFAD gene expression data were downloaded from the NCBI GEO database (accession ID: GSE150358) (`https://www.ncbi.nlm.nih.gov/geo/query/acc.cgi?acc=GSE150358`), originally published in [19].

#### S2 Additional details on GeoAdvAE

##### S2.1 Additional details on architecture

Below, we provide additional implementation details of the GeoAdvAE architecture and training configuration. The model uses a latent-space dimension of $d$=16, with hidden-layer sizes of $\{256, 512, 1024\}$ depending on the modality. All networks were initialized with Kaiming normal initialization and optimized using the Adam optimizer with a batch size of 32 over the total number of iterations. Dropout was applied at a rate of 0.1 in the encoder and decoder networks and 0.3 in the discriminator to prevent overfitting. The prior-guided loss employed a temperature parameter of $\tau$=0.07, while label smoothing with $\epsilon$=0.1 was used in the adversarial discriminator to stabilize training. For the Gromov-Wasserstein (GW) alignment, we used the global alignment setting across all batches. When estimating cell-type proportions during evaluation, the neighborhood size was set to a maximum of $k$=10 for nearest-neighbor computations.

##### S2.2 Additional details on loss function

Below, we provide additional details about the various terms in GeoAdvAE's loss function.

- **Reconstruction loss**: Each modality has its own decoder $D_m$ and reconstruction $\widehat{\mathbf{x}}_m = D_m(\mathbf{z}_m)$ with $\mathbf{z}_m \sim q_m(\mathbf{z}_m \mid \mathbf{x}_m)$. We use an $L_1$ reconstruction loss,

$$\mathcal{L}_{\text{recon}}^{(m)} = \frac{1}{N_m} \sum_{i=1}^{N_m} \left\| \mathbf{x}_m^{(i)} - \widehat{\mathbf{x}}_m^{(i)} \right\|_1, \qquad \mathcal{L}_{\text{recon}} = \mathcal{L}_{\text{recon}}^{(A)} + \mathcal{L}_{\text{recon}}^{(B)}. \tag{S1}$$

  This $L_1$ loss was also used in Crossmodal-AE [21].
- **KL divergence**: For each modality $m \in \{A, B\}$, the encoder $E_m$ outputs parameters of a diagonal Gaussian $q_m(\mathbf{z}_m \mid \mathbf{x}_m) = \mathcal{N}(\boldsymbol{\mu}_m, \operatorname{diag}(\boldsymbol{\sigma}_m^2))$, with latent dimension $d$. We penalize the KL divergence to the standard normal prior $p(\mathbf{z}) = \mathcal{N}(\mathbf{0}, \mathbf{I})$:

$$\mathcal{L}_{\text{KL}}^{(m)} = \frac{1}{d} D_{\text{KL}}(q_m(\mathbf{z}_m \mid \mathbf{x}_m) \parallel p(\mathbf{z})) = -\frac{1}{2d} \sum_{i=1}^{d} \left(1 + \log \sigma_{m,i}^2 - \mu_{m,i}^2 - \sigma_{m,i}^2\right). \tag{S2}$$

  The total KL loss adds the two modalities:

$$\mathcal{L}_{\text{KL}} = \mathcal{L}_{\text{KL}}^{(A)} + \mathcal{L}_{\text{KL}}^{(B)}. \tag{S3}$$

- **Adversarial classification**: For the alternating training between the generator and discriminator, we adopt an unbalanced alternating strategy in which the discriminator is updated twice for every generator update. This stabilizes adversarial learning and prevents model collapse. Each iteration processes one batch of 32 samples. For example, on the patch-seq dataset (645 cells, 3000 iterations), this corresponds to roughly 150 epochs in total.
- **Prior-guided cluster alignment**: To stabilize early optimization, a linear warmup schedule gradually increases the influence of this loss during the first $T_{\text{warm}}$ iterations:

$$\mathcal{L}_{\text{prior}}^{\text{final}} = \min\left(1, \frac{t}{T_{\text{warm}}}\right) \mathcal{L}_{\text{prior}}, \qquad t = 1, \ldots, T_{\text{warm}}. \tag{S4}$$

##### S2.3 Additional details on training and timing

Training follows a three-stage warm-start activation schedule designed to stabilize optimization:

1. **Phase 1 (0–200 iterations)**: Train both VAEs with $\mathcal{L}_{\text{KL}}$ and $\mathcal{L}_{\text{recon}}$ active, establishing modality-faithful encoders. The prior loss $\mathcal{L}_{\text{prior}}$ is gradually warmed up to introduce coarse semantic alignment.
2. **Phase 2 (200–600 iterations)**: Introduce the adversarial term $\mathcal{L}_{\text{GAN}}$ to align latent marginal distributions while preserving reconstruction accuracy. The discriminator is updated twice per iteration ($n_D$=2) for every encoder-decoder update.
3. **Phase 3 (600–1000 iterations)**: Introduce the structural Gromov-Wasserstein loss $\mathcal{L}_{\text{GW}}$ (either global or groupwise mode) to refine geometric alignment of latent spaces.

Each encoder-decoder pair is optimized with the Adam optimizer (lr $= 2 \times 10^{-4}$, $\beta_1 = 0.5$, $\beta_2 = 0.999$) and a small weight decay ($10^{-4}$). The discriminator uses the same learning rate but maintains independent optimizer states. A step-decay policy (decay factor 0.8 every 800 iterations) is applied to the learning rate to ensure stable convergence across training phases.

*Hyperparameters and default configuration.* GeoAdvAE has multiple hyperparameters, as alluded to in the main text. Empirically, the most sensitive ones are:

- The prior loss weight $\lambda_{\text{prior}}$ and temperature $\tau$, which strongly influence semantic cluster alignment;
- The relative ratio between the reconstruction and GAN hyperparameters, $\lambda_{\text{recon}}$ and $\lambda_{\text{GAN}}$, which controls the balance between information preservation and modality matching;
- The latent dimension $d$, which affects both alignment granularity and stability.

For the remaining parameters, moderate settings ($\lambda_{\text{prior}}$=1.0, $\tau$=0.07, $d$=16) provide the most stable and reproducible convergence.

*Model efficiency and computational time.* GeoAdvAE demonstrates high computational efficiency across datasets. Training on the synthetic simulation dataset completes in approximately 5 minutes for 2000 iterations. For the patch-seq dataset, 3000 iterations require about 20 minutes. On the larger 5xFAD dataset, the model converges within 40 minutes over 9000 iterations, confirming the scalability and practicality of the framework for diverse single-cell integration tasks. The following table summarizes the dataset dimensions and the approximate time required to achieve convergence.

**Table S1. Computational Performance Summary.** Overview of the number of cells, highly variable genes (HVGs), and total training time for each analysis. Times are approximate based on standard hardware configurations.

| **Experiment** | **# of Cells** | **# of HVG Genes** | **Training Time** |
|---|---|---|---|
| Simulation | 1,000 | 3 | ≈ 5 min |
| patch-seq (645 cells) | 645 | 2,000 | ≈ 20 min |
| patch-seq (Imbalanced) | 645 Morph / 1329 GEX | 42,466 | ≈ 1h 15min |
| Microglia Analysis | 98 Morph / 31,948 GEX | 2,008 | ≈ 40 min |

As shown in Table S1, GeoAdvAE demonstrates high efficiency even in settings with extreme modality imbalance or high feature counts. This scalability confirms that our framework is viable for large-scale single-cell atlases where joint profiling is impractical.

### S3 Additional details on analysis workflow

#### S3.1 Additional details on simulating data

**Three neuron type simulation.** To systematically validate GeoAdvAE, we designed a biologically interpretable simulation framework that jointly generates synthetic gene expression (GEX) and morphological data with known cross-modal relationships. The simulation is constructed to emulate three canonical neuron archetypes (pyramidal, multipolar, and bipolar neurons), each characterized by distinct transcriptional programs and morphological phenotypes.

The overall generative process follows the chain:

$$\text{Gene expression } (g_1, g_2, g_3) \xrightarrow{\text{sigmoid mapping}} (P, D, A) \xrightarrow{\text{morphology rules}} \text{Neuronal morphology.}$$

This design ensures transparency and interpretability (each gene regulates one distinct morphological axis), separability (distinct clusters form in both modalities), and determinism (identical expression profiles yield identical morphologies), thereby providing a rigorous ground-truth benchmark for evaluating GeoAdvAE's diagonal integration performance.

*Step 1. Generating separable gene expression clusters.* We define three well-separated transcriptional clusters, each corresponding to a distinct neuronal subtype, by sampling three synthetic genes $(g_1, g_2, g_3)$ from Gaussian distributions with distinct means and low variance:

$$\text{Cluster 1 (Pyramidal)}: g_1 \sim \mathcal{N}(2.0,\, 0.2^2), \quad g_2 \sim \mathcal{N}(0.0,\, 0.2^2), \quad g_3 \sim \mathcal{N}(1.2,\, 0.2^2),$$

$$\text{Cluster 2 (Multipolar)}: g_1 \sim \mathcal{N}(-2.0,\, 0.2^2), \quad g_2 \sim \mathcal{N}(2.0,\, 0.2^2), \quad g_3 \sim \mathcal{N}(-1.0,\, 0.2^2),$$

$$\text{Cluster 3 (Bipolar)}: g_1 \sim \mathcal{N}(0.0,\, 0.2^2), \quad g_2 \sim \mathcal{N}(-1.0,\, 0.2^2), \quad g_3 \sim \mathcal{N}(2.0,\, 0.2^2).$$

The resulting clusters are linearly separable in the 3D gene-expression space, establishing an interpretable transcriptional ground truth for downstream alignment analysis.

*Step 2. Mapping GEX to morphological control parameters.* Each gene expression vector $(g_1, g_2, g_3)$ is transformed into three biologically interpretable morphological control parameters (polarity $P$, proximal branching density $D$, and anisotropy $A$) via a nonlinear sigmoid mapping:

$$\sigma(x) = \frac{1}{1 + e^{-x}}, \qquad P = \sigma(1.5 \cdot g_1),\ D = \sigma(1.5 \cdot g_2),\ A = \sigma(1.5 \cdot g_3). \tag{S5}$$

This mapping captures how specific transcriptional programs can regulate macro-scale morphological features. For instance, pyramidal neurons (high $P$, moderate $D$, high $A$) are simulated with a dominant apical trunk and prominent distal branch clusters; multipolar neurons (low $P$, high $D$, low $A$) exhibit dense, isotropic branching; and bipolar neurons (intermediate $P$, low $D$, high $A$) form elongated bidirectional projections.

*Step 3. Morphology generation via procedural rules.* Given the control parameters $(P, D, A)$, we procedurally generate synthetic neuronal trees reflecting their structural archetypes:

- **Pyramidal ($\overline{P}$=0.95, $\overline{D}$=0.50, $\overline{A}$=0.87)**: A single apical trunk extending along the $y$-axis, branching into three major apical clusters alongside 6 basal dendrites; the morphology `.swc` file for each neuron consists of ~200–300 points.
- **Multipolar ($\overline{P}$=0.05, $\overline{D}$=0.95, $\overline{A}$=0.18)**: Approximately 10 primary dendrites radiating isotropically with a branching probability of $p_b = 0.7e^{-d/2}$, forming compact bushy structures; the morphology `.swc` file for each neuron consists of ~150–250 nodes.
- **Bipolar ($\overline{P}$=0.50, $\overline{D}$=0.18, $\overline{A}$=0.95)**: Two symmetric trunks extending along the $\pm y$-axis, each forming distal trifurcations with minimal lateral branching; the morphology `.swc` file for each neuron consists of ~100–200 nodes.

The reported values $\overline{P}$, $\overline{D}$, and $\overline{A}$ denote the empirical means derived from the sampled $(g_1, g_2, g_3)$ coordinates. These rule-based generative procedures ensure that structural geometry remains deterministic and directly interpretable given the underlying gene expression.

*Step 4. Embedding and normalization.* All generated morphologies were embedded into a fixed-length feature representation using the CAJAL framework [8], which computes pairwise Wasserstein distances between neuronal skeletons and derives low-dimensional morphology embeddings.

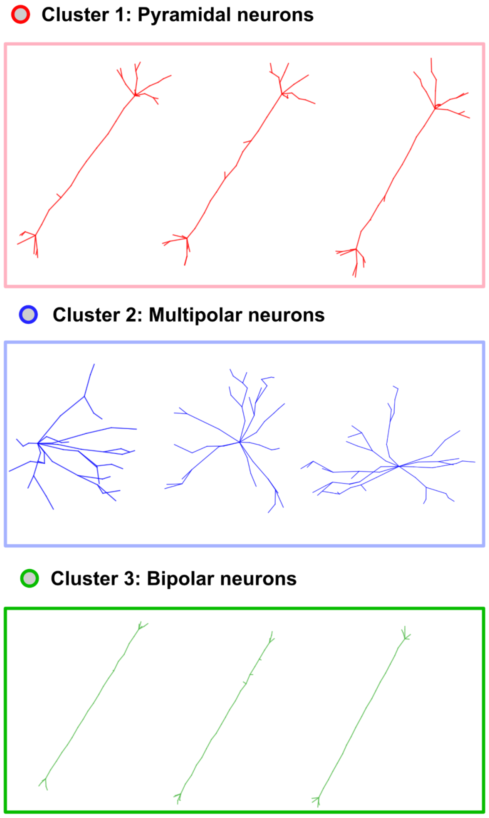


**Fig. S1. Example simulated morphologies for the 3-cluster framework.** Representative simulated neuron morphologies from three predefined clusters: Cluster 1 (red) — pyramidal neurons; Cluster 2 (blue) — multipolar neurons; Cluster 3 (green) — bipolar neurons. These synthetic geometries mimic diverse neuronal arborization patterns for controlled simulation benchmarks.

**Five neuron type simulation** To evaluate GeoAdvAE under a more challenging and realistic scenario, we designed a multi-level hierarchical simulation framework. In real-world single-modality studies, researchers typically have access to broad cell-type labels (e.g., excitatory vs. inhibitory neurons) from canonical marker genes, whereas finer sub-type boundaries must be discovered entirely unsupervised. To emulate this setting, we simulated five distinct fine-grained neuronal subclasses (PA, PB, SA, SB, BI) nested within three coarse main classes: Pyramidal (PA, PB), Stellate (SA, SB), and Bipolar (BI).

The coarse three-class correspondence matrix $\mathbf{P}$ is provided to GeoAdvAE as a structural prior, while the true five-subclass structure is completely withheld from the model during training and reserved exclusively for downstream evaluation. Conditioned on the subclass label, both modalities are generated independently, enforcing a strictly diagonal, unpaired integration task that specifically tests whether the model's geometry-preservation mechanism can resolve hidden boundaries.

The simulation workflow utilizes an expanded five-dimensional version of the process described in the 3-cluster framework, mapping five synthetic genes ($g_1, \ldots, g_5$) to distinct morphological axes: apical polarity, compactness, branching density, vertical elongation, and radial symmetry.

*Step 1. Generating gene expression profiles with asymmetric noise.* For each cell of subclass $s$, the 5D expression vector is drawn from a diagonal Gaussian distribution $\mathbf{g} \sim \mathcal{N}(\boldsymbol{\mu}_s, \sigma_s^2 \mathbf{I}_5)$ parameterized by the following subclass means and noise levels:

$$\begin{aligned}
\text{PA (Tall Pyramidal)} &: \boldsymbol{\mu}_{\text{PA}} = (3.0,\ -2.0,\ -0.5,\ 3.5,\ -2.5), \quad \sigma_{\text{PA}} = 0.25,\\
\text{PB (Wide Pyramidal)} &: \boldsymbol{\mu}_{\text{PB}} = (2.5,\ -1.5,\ 2.0,\ 0.5,\ -2.0), \quad \sigma_{\text{PB}} = 0.25,\\
\text{SA (Dense Stellate)} &: \boldsymbol{\mu}_{\text{SA}} = (-2.0,\ 1.5,\ 1.5,\ -1.0,\ 3.0), \quad \sigma_{\text{SA}} = 0.35,\\
\text{SB (Sparse Stellate)} &: \boldsymbol{\mu}_{\text{SB}} = (-2.5,\ 0.3,\ -0.3,\ 0.5,\ 2.5), \quad \sigma_{\text{SB}} = 0.35,\\
\text{BI (Bipolar)} &: \boldsymbol{\mu}_{\text{BI}} = (0.0,\ -2.5,\ -2.5,\ 4.0,\ -1.0), \quad \sigma_{\text{BI}} = 0.25.
\end{aligned}$$

To thoroughly test the model, we engineered an informational asymmetry into the dataset. While the two Pyramidal sub-types retain large transcriptomic separation, the Stellate subtypes (SA and SB) are positioned intentionally close in GEX space ($\|\boldsymbol{\mu}_{\text{SA}} - \boldsymbol{\mu}_{\text{SB}}\|_2 \approx 3.0$) with elevated noise ($\sigma = 0.35$). Consequently, transcriptomic evidence alone is insufficient to distinguish the two subclasses, requiring the model to leverage morphological geometry to resolve the true biological boundaries.

*Step 2. Procedural morphology generation.* Conditioned on the subclass label, each neuron is procedurally grown into a 3D branching tree and saved in SWC format using the iterative segment extension and stochastic branch-spawning mechanics outlined in the 3-cluster framework. The five subclasses are parameterized to form geometrically extreme, non-overlapping morphological archetypes (visualized in Fig. S2):

- **PA (Tall Pyramidal)**: Characterized by a single, straight apical trunk ($L \sim \mathcal{U}(150, 200)\,\mu$m) terminating in a sparse tuft and compact basal dendrites, exhibiting extreme vertical elongation with a narrow lateral footprint.
- **PB (Wide Pyramidal)**: Characterized by a short apical trunk terminating in a recursively expanding, wide-spreading branched tuft and horizontal basal dendrites, producing a broad mushroom-like shape.
- **SA (Dense Stellate)**: Generated via a breadth-first search (BFS) tree along 20 primary directions uniformly placed over a sphere via a Fibonacci lattice, forming a compact spherical pom-pom with dense, isotropic branching.
- **SB (Sparse Stellate)**: Consists of 5 primary dendrites distributed evenly in azimuth that grow as straight radial chains with minimal branching mass, forming a sparse starfish morphology with large radial reach.
- **BI (Bipolar)**: Consists of two nearly straight opposing trunks projecting along the $\pm y$-axis capped by short terminal sprouts with no lateral dendrites, exhibiting a narrow cigar/rod shape.

In total, 200 cells are generated per subclass ($N = 1,000$ total), yielding 1,000 SWC morphology files and 1,000 completely unpaired five-dimensional gene expression vectors.

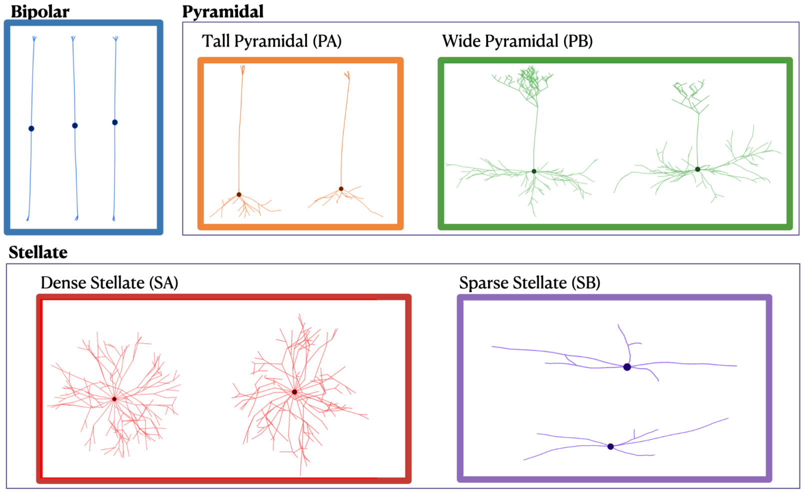


**Fig. S2. Example simulated morphologies for the 5-cluster framework.** Representative simulated neuron morphologies from the five predefined subclasses spanning three main classes: PA (Tall Pyramidal), PB (Wide Pyramidal), SA (Dense Stellate), SB (Sparse Stellate), and BI (Bipolar).

*Step 3. Pre-integration distribution and latent alignment results.* The generated morphologies were vectorized via the identical CAJAL framework described in Step 4 of the 3-cluster framework. Prior to running multi-modal integration, the leading principal components of each modality reveal completely distinct coordinate spaces, unaligned geometric orientations, and unequal scales (Fig. S3). In the independent transcriptomic space, the engineered informational asymmetry is clearly visible: the Dense Stellate (SA) and Sparse Stellate (SB) subclasses heavily blend together due to high coordinate noise, making them indistinguishable via single-modality cluster detection. Conversely, the morphology-derived CAJAL embeddings exhibit structural separability but remain entirely unmapped to the GEX profile vectors.

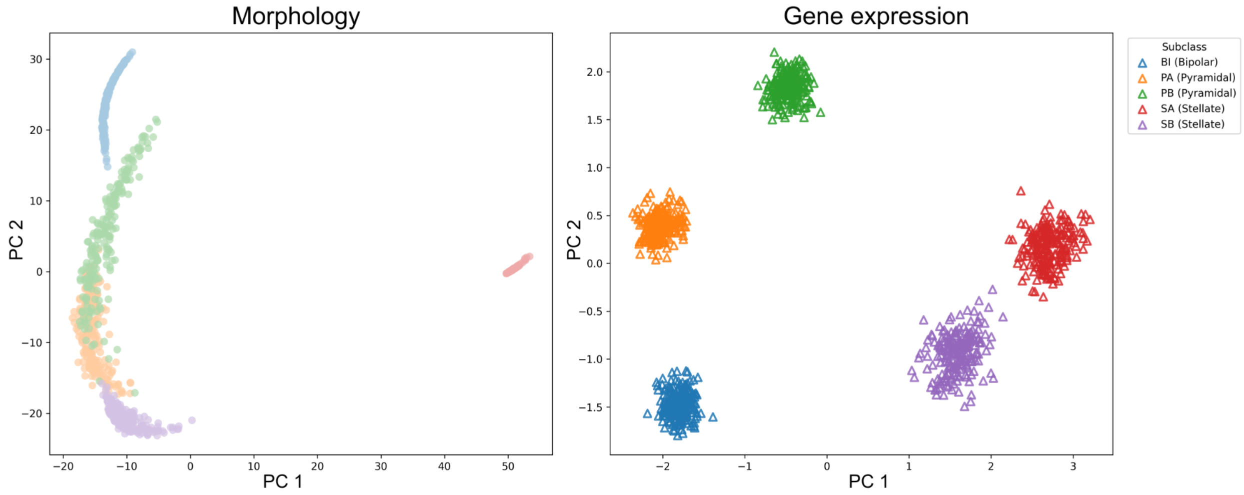


**Fig. S3. Pre-alignment distributions of the 5-subclass simulation in independent PCA spaces.** Projections of the unaligned single-modality datasets onto their respective leading principal components prior to integration. In the gene expression space (triangles), the Stellate subtypes (SA and SB) exhibit high overlap and blending due to the engineered coordinate noise. In the morphological space (filled circles), CAJAL embeddings resolve distinct structural geometry but occupy an entirely separate feature space with no explicit matching or coordinate alignment to the transcriptomic profiles, establishing a rigorous validation case for geometry-preserving diagonal integration.

Following diagonal integration with GeoAdvAE, the joint cross-modal latent space was projected into PCA space to verify alignment quality. As shown in Fig. S4, GeoAdvAE achieved near-perfect multi-modal alignment across all hidden sub-classes. Even though the coarse prior matrix $\mathbf{P}$ groups SA and SB together into a single broad "Stellate" category and transcriptomic signals blur between them, GeoAdvAE's Gromov-Wasserstein loss successfully uncovered the withheld SA/SB sub-class boundary by preserving morphological geometry. The final integrated space achieved a cross-modal 1-nearest-neighbor (1NN) cell-type matching accuracy of 0.99, confirming that our geometry-aware regularization can accurately resolve granular structures that are completely omitted from coarse biological priors.

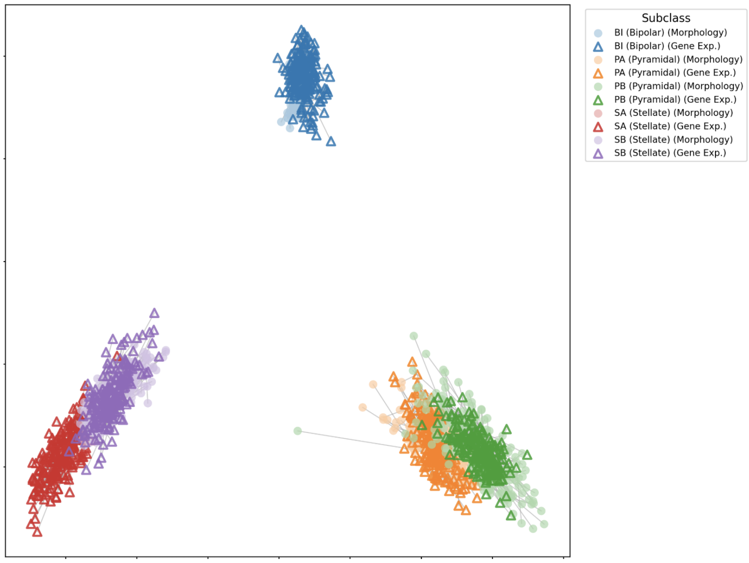


**Fig. S4. Cross-modal alignment of simulated neuron dataset in PCA space.** Principal component analysis representation of the integrated multi-modal dataset. Filled circles represent morphology-derived embeddings, and triangles indicate gene expression profiles. Cohesive clustering and substantial spatial overlap within each hidden subclass validate successful alignment, achieving a 1NN cell-type matching accuracy of 0.99 under coarse prior guidance.

#### S3.2 Additional details on cluster alignment loss for patch-seq data

To construct the correspondence matrix $\mathbf{P}$ mapping broad clusters in GEX to morphology used in the prior-guided alignment penalty for the patch-seq dataset, we incorporated biological prior knowledge about excitatory-inhibitory (E/I) cell type correspondence between the gene expression (GEX) and morphology modalities. The rationale behind this design is that, although the two modalities are unpaired at the single-cell level, excitatory and inhibitory neurons exhibit consistent molecular and structural characteristics across modalities.

For the GEX modality, we clustered cells into 4 clusters. Each cluster was annotated as excitatory (E) or inhibitory (I) based on canonical marker genes stated in [11]. This yielded the GEX-type proportion matrix:

$$\pi_{\text{GEX}} = \begin{array}{c|cccc} & \text{Cluster 1} & \text{Cluster 2} & \text{Cluster 3} & \text{Cluster 4} \\ \hline \text{E} & 0 & 1 & 0 & 0 \\ \text{I} & 1 & 0 & 1 & 1 \end{array}.$$

Here, each element $\pi_{\text{GEX}}(t, i)$ represents the proportion of cells of type $t \in \{\text{E}, \text{I}\}$ within GEX cluster $i$.

For the morphology modality, we sampled 10 representative neuron morphologies from each cluster and manually classified them as excitatory or inhibitory based on dendritic and axonal branching patterns via visual inspection. This yielded the morphology-type proportion matrix:

$$\pi_{\text{Morpho}} = \begin{array}{c|cccc} & \text{Cluster 1} & \text{Cluster 2} & \text{Cluster 3} & \text{Cluster 4} \\ \hline \text{E} & 0.2 & 0.7 & 0.1 & 0.9 \\ \text{I} & 0.8 & 0.3 & 0.0 & 0.1 \end{array}.$$

Each element $\pi_{\text{Morpho}}(t, j)$ denotes the proportion of cells of type $t$ in morphology cluster $j$, obtained from manual inspection.

Because excitatory and inhibitory neurons are expected to correspond across modalities, i.e., excitatory morphologies align to excitatory GEX profiles and inhibitory morphologies align to inhibitory ones, we defined the following intermediary matrix:

$$\mathbf{R} = \begin{array}{c|cc} & \text{E} & \text{I} \\ \hline \text{E} & 1 & 0 \\ \text{I} & 0 & 1 \end{array}.$$

The final correspondence matrix $\mathbf{P}$ was then computed as

$$\mathbf{P} = \pi_{\text{GEX}}^{\top} \cdot \mathbf{R} \cdot \pi_{\text{Morpho}},$$

where each entry $P_{ij}$ quantifies the biological correspondence between GEX cluster $i$ and morphology cluster $j$ based on their shared excitatory-inhibitory composition.

The resulting correspondence matrix $\mathbf{P}$ is shown below:

$\mathbf{P} =$

| | Morpho Cluster 1 | Morpho Cluster 2 | Morpho Cluster 3 | Morpho Cluster 4 |
|---|---|---|---|---|
| GEX Cluster 1 | 0.80 | 0.30 | 0.90 | 0.10 |
| GEX Cluster 2 | 0.15 | 0.70 | 0.10 | 1.00 |
| GEX Cluster 3 | 0.80 | 0.30 | 0.90 | 0.10 |
| GEX Cluster 4 | 0.75 | 0.25 | 0.85 | 0.05 |

Rows correspond to GEX clusters ($i$), and columns correspond to morphology clusters ($j$). Each element $P_{ij}$ represents the prior correspondence strength between GEX cluster $i$ and morphology cluster $j$, derived from the shared excitatory-inhibitory composition across modalities. This matrix was used during GeoAdvAE training.

##### S3.3 Additional details on cluster alignment loss for 5xFAD data

We document our procedure for constructing the correspondence matrix $\mathbf{P}$ for the 5xFAD microglia analysis, which differs slightly from our patch-seq analysis due to a more "smooth" continuum among different microglial states.

*Morphology prior.* For the morphology modality, we clustered the 98 microglia based on the CAJAL embeddings and manually annotated them via visual inspection by broad canonical shapes (cf. Fig. 5): *Ramified*, *Intermediate*, and *Amoeboid*. This yields a morphology-type proportion matrix that is an identity mapping:

$\pi_{\text{Morpho}} =$

| | Ramified | Intermediate | Amoeboid |
|---|---|---|---|
| Ramified | 1 | 0 | 0 |
| Intermediate | 0 | 1 | 0 |
| Amoeboid | 0 | 0 | 1 |

*GEX prior.* The prior matrix for the GEX modality was more complex, since we did not want to represent each microglia cluster, as originally derived by the authors (Wang et al., [19]), as a monolithic cluster. Many studies of microglia have uncovered a continuum spanning disease progression [9,7], suggesting that using "discrete" clusterings to construct our correspondence matrix $\mathbf{P}$ may be suboptimal. Hence, we further refined the clustering by Wang et al. (12 original clusters) by representing each cluster as a collection of distinct microglial states. We score each cluster based on the following marker panel, derived by reviewing the literature [19,9,18]:

Homeostatic : {*P2ry12*, *Tmem119*, *Cx3cr1*}, Proliferating : {*Mki67*, *Top2a*, *Pcna*},

IRM : {*Ifit1*, *Ifit3*, *Isg15*, *Stat1*}, DAM : {*Apoe*, *Trem2*, *Axl*, *Cst7*}.

Using these four panels, we computed the mean expression across markers and cells in each cluster to obtain a raw score per state, and then normalized the four raw scores. This resulted in percentages that sum up to 100 percent for each cluster, which is represented by the $\pi_{\text{GEX}}$ matrix:

$\pi_{\text{GEX}} =$

| | H1 | H2 | H3 | DAM 2 | H4 | H5 | IRM | Transition | DAM 1 | P1 | P2 | P3 |
|---|---|---|---|---|---|---|---|---|---|---|---|---|
| Homeostatic | 88 | 93 | 81 | 36 | 65 | 80 | 62 | 77 | 16 | 47 | 84 | 49 |
| Proliferating | 1 | 1 | 1 | 1 | 1 | 1 | 1 | 1 | 0 | 19 | 2 | 37 |
| IRM | 1 | 1 | 1 | 1 | 1 | 1 | 11 | 2 | 1 | 3 | 1 | 2 |
| DAM | 11 | 6 | 18 | 63 | 33 | 19 | 26 | 20 | 83 | 31 | 13 | 13 |

Each element $\pi_{\text{GEX}}(t, i)$ denotes the percentage of state $t \in$ {Homeostatic, Proliferating, IRM, DAM} within GEX cluster $i$. In this matrix, each row represents the relative enrichment of one transcriptional state, and each column corresponds to one of the twelve GEX clusters (*H1–H5*, *DAM1–2*, *Transition*, *IRM*, *P1–P3*). Among these, *H1–H5* denote five subtypes of homeostatic microglia, while *P1–P3* represent proliferating subclusters. As expected, the homeostatic clusters (*H1–H5*) are predominantly enriched for homeostatic marker genes, whereas the proliferating clusters (*P1–P3*) show higher scores for proliferative markers. Similarly, the clusters *DAM1–2* and *IRM* are characterized by strong enrichment of DAM- and IRM-associated signatures, respectively. Although all values in $\pi_{\text{GEX}}$ are expressed as proportions, the enrichment pattern clearly reflects biologically meaningful state-specific compositions.

*Correspondence matrix.* To derive the correspondence matrix $\mathbf{P}$, we first manually construct the following intermediary matrix $\mathbf{R}$ that maps broad microglia GEX profiles to broad microglia morphologies after reviewing the literature. We specifically used the following $\mathbf{R}$ in our 5xFAD analysis:

$\mathbf{R} =$

| | Ramified | Intermediate | Amoeboid |
|---|---|---|---|
| Homeostatic | 1 | 0.2 | 0 |
| Proliferating | 0.2 | 1 | 0.2 |
| IRM | 0 | 0.4 | 1 |
| DAM | 0 | 0.2 | 1 |

.

The weights that are not strictly 0 or 1 were assessed by reviewing images of microglia in different states [15,10,6,3], where the relative weights were concretely decided based on how much we believe the evidence suggests a particular microglia state reflected a particular morphology. We performed sensitivity analyses to assess that our downstream biological findings did not change based on the precise values in $(0, 1)$ in the entries of $\mathbf{R}$ that were not strictly 0 or 1.

Given $\pi_{\text{GEX}} \in \mathbb{R}^{4\times12}$, $\mathbf{R} \in \mathbb{R}^{4\times3}$, and $\pi_{\text{Morpho}} \in \mathbb{R}^{3\times3}$, the cluster-level prior between GEX cluster $i$ and morphology cluster $j$ is computed by

$$\mathbf{P} \;=\; \pi_{\text{GEX}}^{\top} \cdot \mathbf{R} \cdot \pi_{\text{Morpho}} \;\;\in\; \mathbb{R}^{12\times3},$$

where rows correspond to 12 GEX clusters annotated by Wang et al. and columns to morphology clusters (Ramified, Intermediate, Amoeboid). Each entry $P_{ij}$ reflects the expected correspondence strength derived from $\mathbf{R}$.

##### S3.4 Additional details of GSEA

We perform GSEA using the `clusterProfiler` R package [22]. Specifically, we use the `clusterProfiler::gseGO()` function with `minGSSize=10` and `maxGSSize=500`. We only analyze the biological process ontology (i.e., `ont="BP"`) using the pathways from the `org.Mm.eg.db` (for the mouse). The inputs to GSEA are the `mean_attribution` (importance scores) for each gene, derived from integrated gradients.

##### S3.5 Additional details about comparison methods for diagonal integration

The diagonal integration methods we compare against span a range of original data settings, several of which differ from our unpaired morphology-gene expression problem. First, SCOT [5], MMD-MA [14], UnionCom [2], and SCIM [17] were built for the same regime we study (i.e., two modalities measured in different cells with no shared features and no cell-level pairing) and align modalities through geometry, distribution, or adversarial latent matching without any feature correspondence. Second, scJoint [13], sciCAN [20], and scDART [24] were designed for unpaired scRNA-seq and scATAC-seq integration, where a gene-activity prior places both modalities on a shared gene-level feature axis. Third, Crossmodal-AE [21] integrates single-cell imaging with sequencing measured in different cells through a shared latent space, while STACI [23] was developed for spatial transcriptomics and chromatin imaging that are co-registered in the same tissue and linked by a spatial-neighborhood graph. Finally, CycleGAN [25] originates in computer vision, where it performs unpaired image-to-image translation.

Because cellular morphology and gene expression share no common feature axis and are profiled in different cells, the methods that presuppose such structure cannot be applied to our setting without modification: the scRNA-seq and scATAC-seq methods have no morphological analog of a gene-activity matrix, and

STACI was developed for a spatial, co-registered setting that our dissociated morphology and expression do not share. Hence, we implemented a representative baseline for these methods on a standardized framework using on same CAJAL-encoded morphologies and log-normalized gene expression inputs, where the original gene-activity matrices or spatial-neighborhood graphs were not used. We present only the methods that rely on a gene-activity or spatial prior (scJoint, sciCAN, scDART, STACI) as "-like" baselines, reflecting that we use the integration paradigm rather than apply the original method literally. This way, our simulation and patch-seq studies still span each integration paradigm (i.e., optimal transport, distribution matching, adversarial latent alignment, cycle-consistency, graph autoencoders, and transfer learning embedding), each with at least one representative baseline.

### S4 Additional results

#### S4.1 Additional results on simulation: Shapley value analysis for loss component importance

To provide a rigorous quantitative assessment of each loss term's contribution to the GeoAdvAE objective function, we conducted a systematic ablation study using Shapley values. While the main text demonstrates that the full model achieves 100% accuracy on simulated data, this analysis identifies which components are most critical for cross-modal alignment and orientation.

We utilized a combinatorial approach on our simulated dataset, systematically enabling and disabling the four primary loss components: Reconstruction ($\mathcal{L}_{\text{recon}}$), Adversarial ($\mathcal{L}_{\text{GAN}}$), Prior-guided alignment ($\mathcal{L}_{\text{prior}}$), and Gromov-Wasserstein ($\mathcal{L}_{\text{GW}}$). This resulted in $2^4 - 1 = 15$ distinct model configurations. For each configuration, we measured the performance using the 1-KNN cell-type matching accuracy.

The importance of each loss term $i$ was then computed as its Shapley value ($\phi_i$), which represents the average marginal contribution of that term across all possible subsets of components:

$$\phi_i = \sum_{S \subseteq N \setminus \{i\}} \frac{|S|!(n - |S| - 1)!}{n!} [v(S \cup \{i\}) - v(S)] \tag{S6}$$

where $N$ is the set of all four loss terms, $n = |N|$, $S$ is a subset of terms excluding $i$, and $v(S)$ is the 1-KNN matching accuracy achieved by the configuration using only the terms in $S$.

The results of the 15 configurations are detailed in Table S2, and the relative importance derived from Shapley values is visualized in Fig. S5.

- **Synergy Requirement:** While the full model (Method 15 in Table S2) achieves 100% accuracy, almost no other combination reaches this threshold. This suggests that high-fidelity diagonal integration is a synergistic process requiring all four geometric and semantic constraints.
- **Dominant Components:** The analysis reinforces that the Adversarial term (34.7%) and the Prior term (31.7%) are the most impactful components. This is biologically and computationally intuitive: the adversarial loss aligns the two latent distributions, while the prior provides the necessary orientation to ensure that excitatory and inhibitory clusters map to their correct counterparts.
- **Refinement Terms:** The Reconstruction and GW terms, while possessing lower Shapley values, act as critical regularizers that stabilize the latent manifold and preserve within-modality geometry, respectively.

**Table S2. Ablation Study Results.** Matching accuracy for all 15 combinations of enabled (denoted by 1) or disabled (denoted by 0) loss components on simulated data.

| Method | Reconstruction | Adversarial | Prior | GW | 1-KNN Accuracy |
|---|---|---|---|---|---|
| 1 | 0 | 1 | 1 | 1 | 0.85 |
| 2 | 1 | 0 | 1 | 1 | 0.84 |
| 3 | 1 | 1 | 0 | 1 | 0.84 |
| 4 | 1 | 1 | 1 | 0 | 0.90 |
| 5 | 0 | 0 | 1 | 1 | 0.59 |
| 6 | 0 | 1 | 0 | 1 | 0.52 |
| 7 | 0 | 1 | 1 | 0 | 0.72 |
| 8 | 1 | 0 | 0 | 1 | 0.25 |
| 9 | 1 | 0 | 1 | 0 | 0.86 |
| 10 | 1 | 1 | 0 | 0 | 0.53 |
| 11 | 0 | 0 | 0 | 1 | 0.49 |
| 12 | 0 | 0 | 1 | 0 | 0.50 |
| 13 | 0 | 1 | 0 | 0 | 0.79 |
| 14 | 1 | 0 | 0 | 0 | 0.36 |
| 15 | 1 | 1 | 1 | 1 | 1.00 |

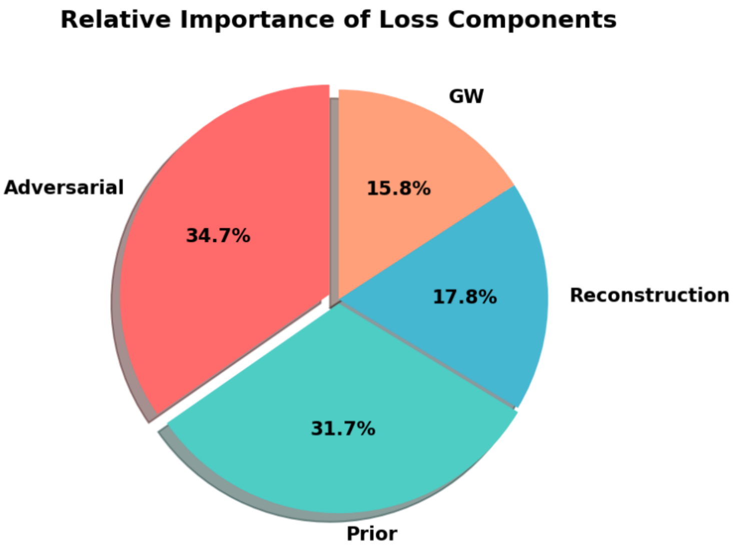


**Fig. S5. Relative Importance of Loss Components.** Shapley value distribution showing that Adversarial and Prior losses contribute over 65% of the model's predictive power for cross-modal alignment.

#### S4.2 Additional results on simulation: Comparison against other diagonal integration methods

Fig. S6 shows the integrated latent spaces of simulated neurons produced by various baseline integration methods. Each point represents a single cell, with circles indicating gene expression embeddings and triangles indicating morphology embeddings; colors correspond to three simulated neuronal clusters. An ideal integration would mix the two modalities within each cluster while maintaining clear separation across clusters. However, most competing methods exhibit fragmented or modality-separated structures, indicating incomplete cross-modal alignment compared to the coherent integration achieved by GeoAdvAE.

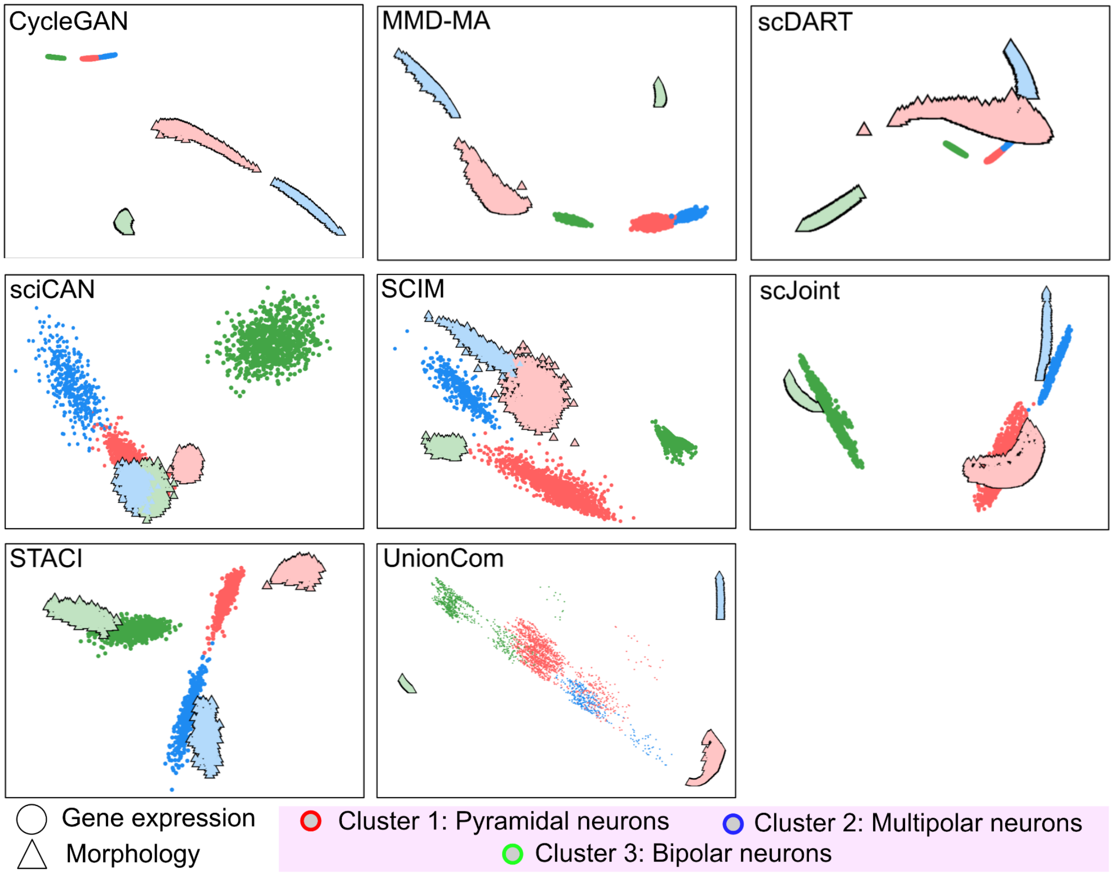


**Fig. S6. Additional results of integration of simulated neurons.** Integrated latent spaces obtained from simulation data using various competing cross-modal integration methods. These plots are shown in the same format as Fig. 2B and 3B.

##### S4.3 Additional results on simulation: Comparison between GW and MMD losses

A central component of the GeoAdvAE framework is the use of the Gromov-Wasserstein (GW) loss to enforce geometric consistency between modalities in the latent space. To justify this choice, we performed a comparative analysis by replacing the GW loss with the Maximum Mean Discrepancy (MMD) loss. This allows us to assess whether a global distribution-matching metric (MMD) is sufficient for diagonal integration or if the relational/intra-modality geometric preservation of GW is necessary for high-fidelity alignment.

*MMD with Laplacian kernel.* We modified the GeoAdvAE loss function by substituting the GW term with an MMD term. We utilized a Laplacian kernel, which is often more robust to outliers in high-dimensional biological data than the standard Gaussian kernel. The MMD squared is calculated as:

$$\mathrm{MMD}^2(Z_a, Z_b) = \frac{1}{n^2}\sum_{i,j} k(z_a^i, z_a^j) + \frac{1}{m^2}\sum_{i,j} k(z_b^i, z_b^j) - \frac{2}{nm}\sum_{i,j} k(z_a^i, z_b^j) \tag{S7}$$

where $k(z, z') = \exp(-\|z - z'\|_1/\sigma)$. We utilized a fixed bandwidth of $\sigma = 1.0$ and the $L_1$ norm (Manhattan distance) for the kernel calculation. We applied this modified loss to the same simulated dataset used in our primary benchmarks and evaluated the resulting latent integration visually and quantitatively via matching accuracy.

*Interpretation of results.* The results of the MMD-based integration are visualized in Fig. S7.

– **Quantitative Performance:** Similar to the GW-based GeoAdvAE, the MMD variant achieves 100% matching accuracy on this simulated dataset, indicating that for simple cluster separations, distribution matching is functionally sufficient.

– **Qualitative Integration:** Despite the high accuracy, visual inspection reveals that the MMD-based integration is less "compact" than the GW-based results. The clusters in the latent space show higher intra-cluster variance and less precise overlap between modalities.

This suggests that while MMD effectively aligns the broad distributions, it lacks the fine-grained geometric constraints provided by GW, which explicitly penalizes the distortion of inter-sample distances within each modality. Consequently, we maintain that GW is the better choice for diagonal integration in complex biological settings where the preservation of manifold structure is paramount.

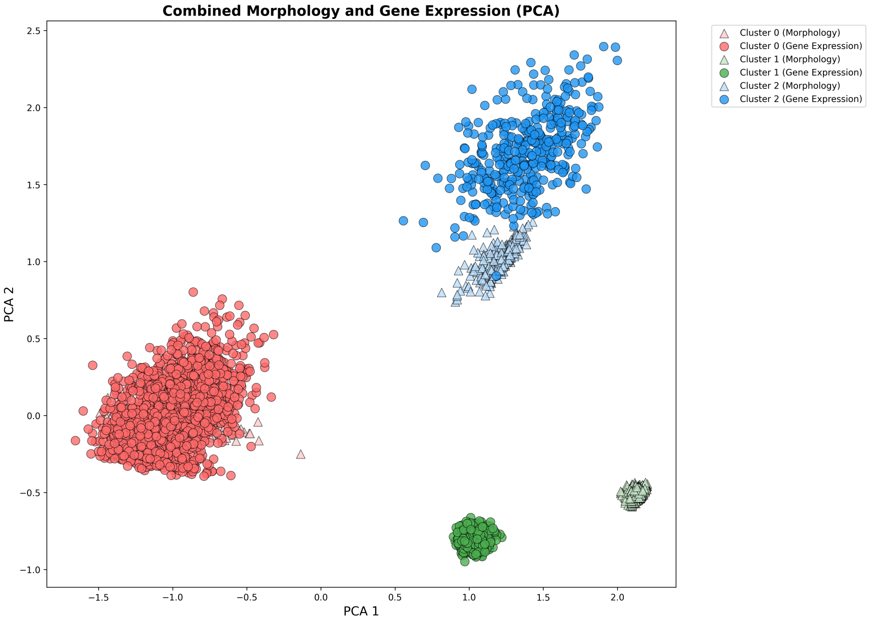


**Fig. S7. Latent Integration via MMD Loss.** Visualizing the shared latent space when replacing GW loss with MMD (Laplacian kernel, $\sigma = 1$). While accuracy remains high, the modalities exhibit less precise geometric overlap compared to the GW-based alignment.

#### S4.4 Additional results on patch-seq: Comparison against other diagonal integration methods

Fig. S8 shows the integrated latent spaces of patch-seq neurons produced by various baseline cross-modal integration methods. Each point represents a single cell, with circles denoting gene expression profiles and triangles denoting corresponding morphological representations. Ideally, cells of the same type should form overlapping, continuous clusters across modalities, indicating successful alignment. However, most competing methods yield fragmented or poorly mixed structures, highlighting the difficulty of achieving coherent cross-modal integration compared with GeoAdvAE.

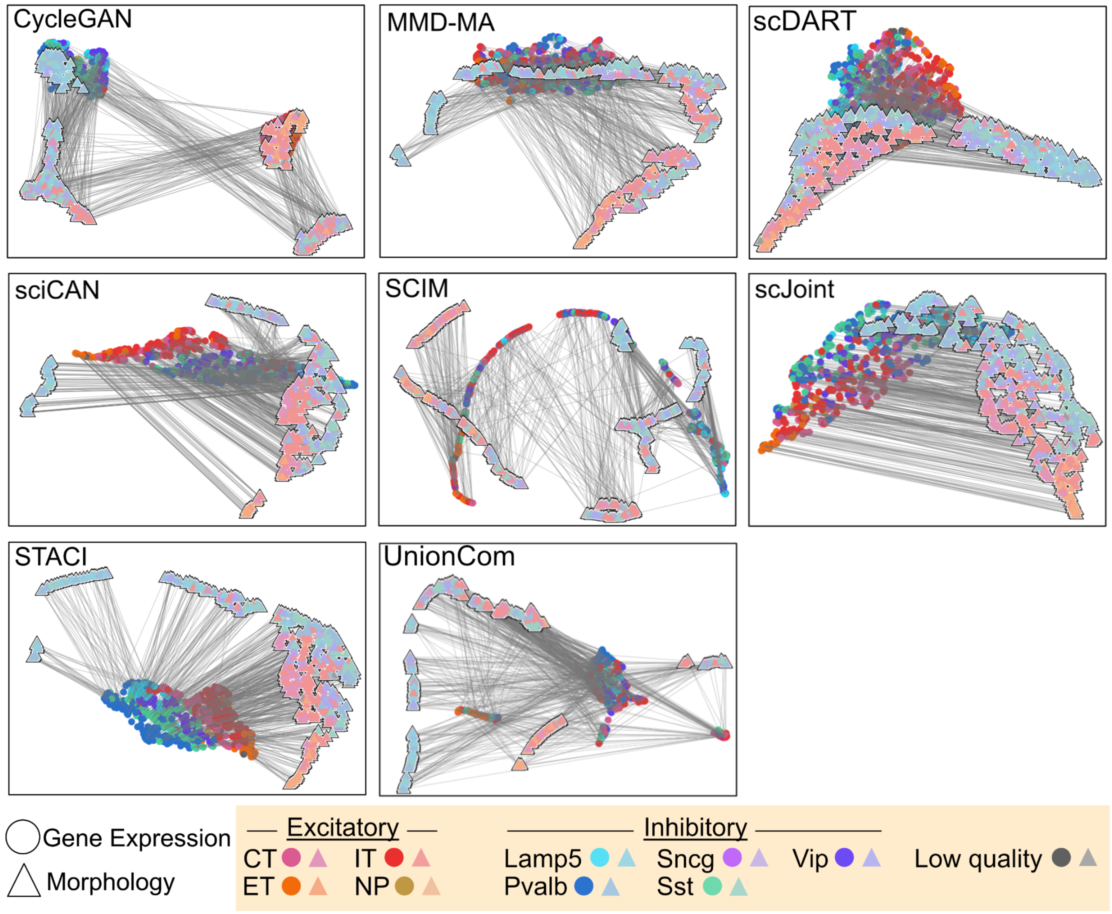


**Fig. S8. Additional results of integration of patch-seq neurons.** Integrated latent spaces obtained from patch-seq data using various competing cross-modal integration methods. These plots are shown in the same format as Fig. 4B,D.

#### S4.5 Additional results on patch-seq: Comparison between Laplacian scores and importance scores

To validate the biological relevance of the genes identified by GeoAdvAE's integrated gradient (IG) attribution, we compared our importance scores against the Laplacian score, a paired graph-based metric for feature selection. This analysis serves as a power analysis demonstrating that our unpaired integration method recovers gene-morphology associations similar to those identified with ground-truth pairings.

The Laplacian score is implemented via the CAJAL framework to assess whether a numerical feature (e.g., gene expression) is related to cellular morphology. This is done using the `cajal.laplacian_scores()` function. Specifically, this calculation involves:

1. **Graph Construction:** An undirected graph $G$ is built where nodes represent cells and edges connect cells with similar morphologies, determined by the Gromov-Wasserstein distance.
2. **Feature Evaluation:** Each gene is passed in as a feature $f$. The Laplacian Score $L_r$ of a gene with respect to the graph $G$ is defined as:

$$L_r = \frac{\sum_{i,j}(f_i - f_j)^2 G_{ij}}{\mathrm{Var}(f)}. \tag{S8}$$

3. **Interpretation:** A low Laplacian score indicates that the gene's expression is "smooth" relative to the morphological graph—meaning cells with similar shapes have similar expression levels for that gene.

In Fig. S9A, we observe a strong negative correlation (Pearson correlation $\approx -0.48$) between GeoAdvAE importance scores (IG) and Laplacian scores. This inverse relationship is biologically consistent: a **high IG** indicates a gene is highly influential in the model's shared latent embedding, while a low Laplacian score indicates a gene is statistically associated with morphological similarity in paired data. The alignment of these metrics demonstrates that even though GeoAdvAE is an unpaired method, it effectively prioritizes the same transcriptomic anchors that paired statistical tests identify as morphology-associated.

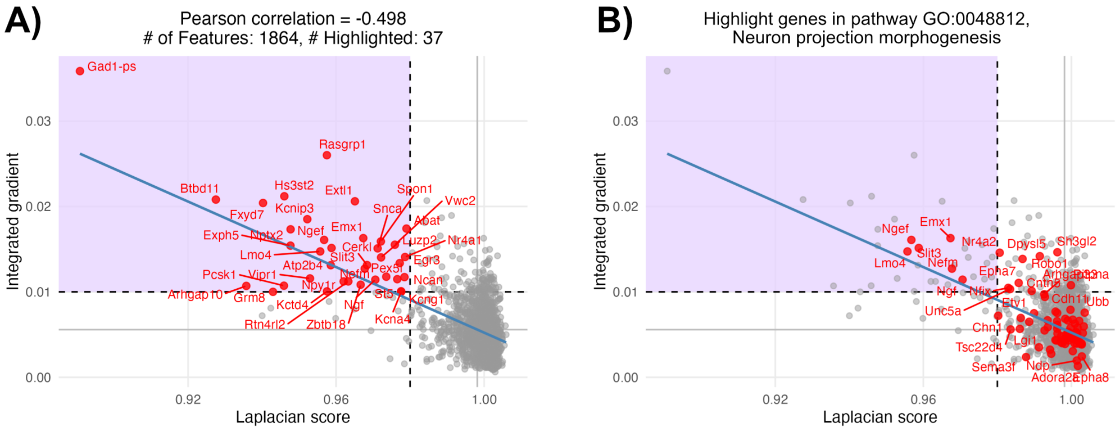


**Fig. S9. Validation of importance scores.** A) Correlation between GeoAdvAE Integrated Gradient scores and CAJAL Laplacian scores, where we annotate genes in red with both a low Laplacian score (high alignment in the paired analysis) and high integrated score (high alignment in the unpaired analysis). B) Same plot, but highlighting genes in red specifically within the Neuron Projection Morphogenesis pathway (GO:0048812).

We highlight several genes that exhibit both high IG and low Laplacian scores, marking them as critical transcriptomic-morphological anchors:

- *Gad1*: Acts as a pan-inhibitory marker, facilitating the separation of GABAergic families that possess largely non-overlapping morpho-electric phenotypes.
- *Rasgrp1*: Essential for the fine-grained classification of excitatory t-types (e.g., ET subclasses) defined by distinct structural features like large apical dendritic tufts.
- Differentiation and Development: Other top-ranked genes include *Lmo4*, *Zbtb18*, *Ngf*, *Slit3*, *Rtn4rl2*, and *Emx1*, all of which are known to play vital roles in neuron differentiation and axon development.

To further demonstrate that GeoAdvAE captures established biological programs, we specifically examined the Neuron Projection Morphogenesis pathway (GO:0048812).

As shown in Fig. S9B, this specific biological pathway is significantly enriched with upregulated genes that possess high IG and low Laplacian scores. The clustering of these genes in the high-importance/high-smoothness quadrant validates that GeoAdvAE identifies a concerted transcriptomic program dedicated to the structural remodeling of the cell. Key highlighted genes within this pathway, such as *Sema3f*, *Epha7*, and *Robo1*, further reinforce the model's ability to nominate candidates that are mechanistically linked to the "form" of the neuron.

#### S4.6 Additional results on patch-seq: Quantifying alignment uncertainty via Monte Carlo transport diagnostics

Mapping high-dimensional gene expression (GEX) to complex morphology is intrinsically challenging. We provide a diagnostic to distinguish robust cross-modal pairings from stochastic noise, identifying which cell states are aligned with high confidence.

*Monte Carlo procedure.* We utilize a Monte Carlo approach based on the mini-batch Gromov-Wasserstein transport plan $T$ estimated during training. This approach is necessitated by the fact that the transportation mapping produced by `ot.gromov.gromov_wasserstein2()` often yields a binary-like (sparse) matrix within a single mini-batch, which does not inherently capture the continuous uncertainty of the global alignment.

The procedure is as follows:

1. **Frequency Estimation:** Over 500 iterations, we sample mini-batches of 32 cells and record the transportation matrix $T$. We aggregate these into a frequency matrix $F$, where $F_{ij}$ counts how often GEX cell $i$ is paired with Morph cell $j$.
2. **Spectral Analysis:** We apply SVD to $F$ to identify latent factors. The singular value spectrum of the frequency matrix is relatively flat (Fig. S10), which is the expected behavior in this context; in an ideal matching scenario where the frequency matrix approaches a diagonal identity, the spectrum would be perfectly flat. We identified $k = 9$ factors as a heuristic elbow to capture the primary alignment structure.
3. **Stochastic Projection:** We average mappings between clusters and project the result into the space of doubly stochastic matrices using iterative row and column normalization (Sinkhorn-style iterations):

$$M_{i,j}^{(t+1)} = \text{Norm}_{col}(\text{Norm}_{row}(M_{i,j}^{(t)})) \quad \text{(S9)}$$

This ensures each cluster's mapping probability sums to 1, offering a normalized confidence score.

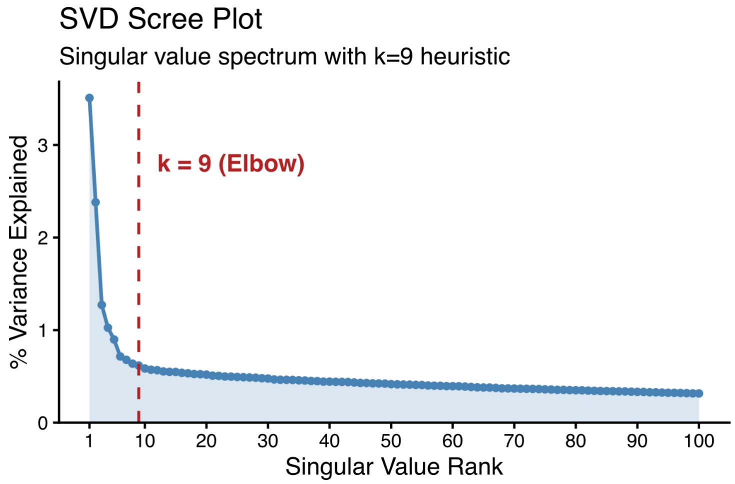


**Fig. S10. SVD scree plot.** Singular value spectrum for the Monte Carlo frequency matrix, showing the $k = 9$ elbow heuristic.

*Interpretation of alignment results.* The results of this diagnostic are visualized in Fig. S11.

- **Type Consistency:** Panel A demonstrates that the alignment primarily respects broad biological categories; excitatory (E) and inhibitory (I) GEX clusters map almost exclusively to their corresponding morphological counterparts.
- **Differential Confidence:** The block-diagonal structure in the binarized matrix suggests that certain cell types/states are more confidently and consistently mapped than others.
- **Cluster Fidelity:** Panel B provides a normalized confidence score for inter-cluster mappings. The high values along the diagonal indicate that for most clusters, the model identifies a "dominant" morphological state, though the off-diagonal values provide a quantifiable measure of mapping ambiguity or biological "fuzziness" between states.

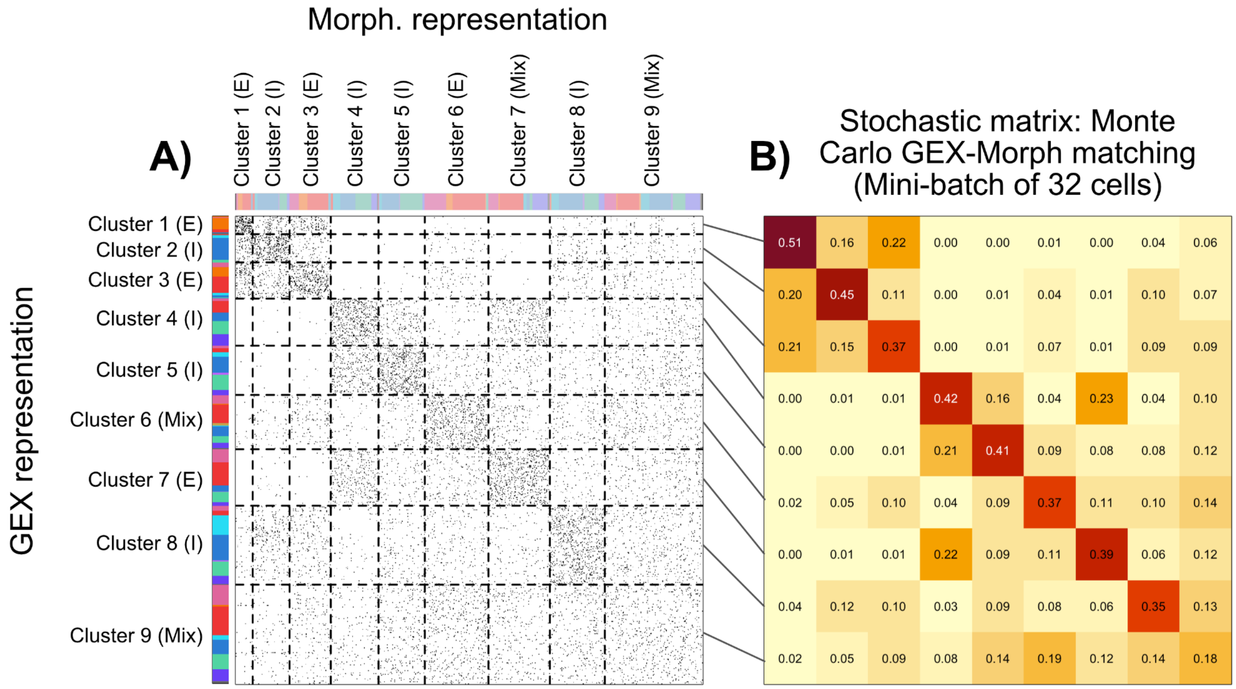


**Fig. S11. Cross-modal mapping diagnostics.** A) Binarized frequency matrix of cell pairings grouped by SVD clusters. B) Doubly stochastic matrix representing normalized mapping confidence between GEX and morphological clusters. The same ordering of clusters (rows: GEX; columns: Morphology) is preserved from (A).

##### S4.7 Additional results on patch-seq: Comparative analysis of CAJAL versus classical morphological features

One finding of our work is that metric geometry-based descriptors, as provided by the CAJAL framework, offer a more holistic representation of cellular morphology than traditionally measured discrete features. To test this, we compared the integration performance of GeoAdvAE using CAJAL embeddings with that of a set of classically measured neuronal morphological features.

*Procedure and feature normalization* We utilized a comprehensive set of morphological features extracted from the patch-seq dataset (e.g., total length, number of branch points, max Euclidean distance, and tortuosity for both apical dendrites and axons). Because certain features (such as apical-specific measurements) are not consistently measured across all excitatory and inhibitory neurons, we applied the following normalization procedure:

1. **Scaling:** All measurements for a specific feature were scaled to the interval $[1, 2]$ based on the observed maximum and minimum values.
2. **Imputation:** Missing values were set to 0 to distinguish them from measured values.
3. **Training:** These normalized classical features were used as the input modality for GeoAdvAE, replacing the CAJAL embeddings, while keeping all other hyperparameters constant.

*Interpretation of results.* The resulting integrated latent space is visualized in Fig. S12. Quantitatively, the overall 1-KNN cell-type matching accuracy was 0.25, which is significantly lower than the 0.34 achieved using CAJAL embeddings. Qualitatively, the shared latent space generated with classical features appears less cohesive. Consequently, we maintain that metric geometry representations provide a better basis for linking “form” and “function” in single-cell data.

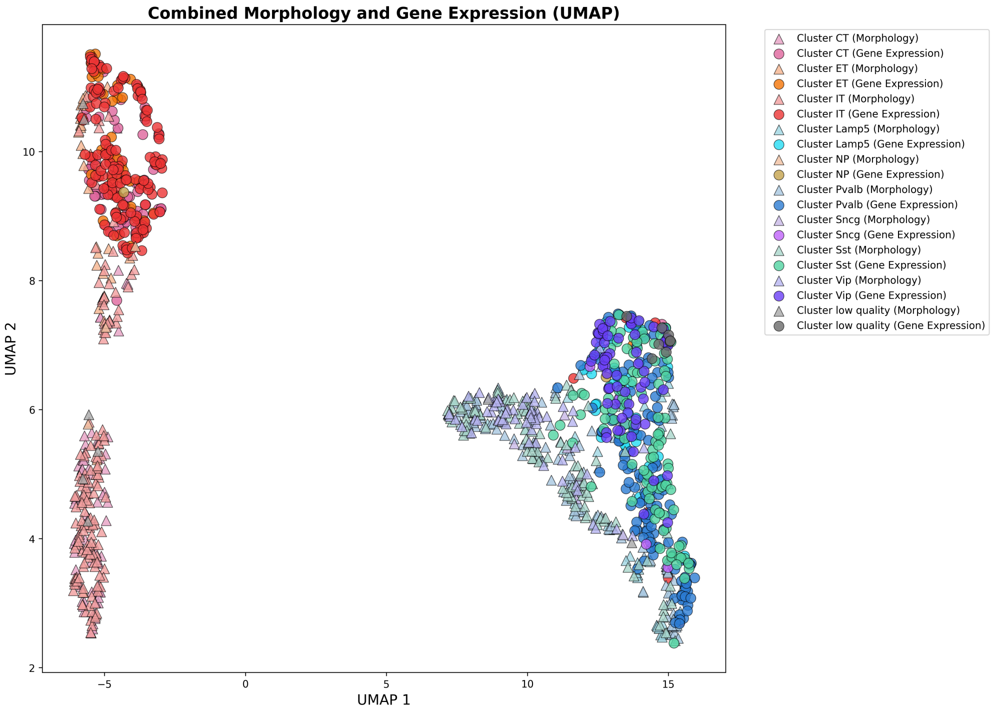


**Fig. S12. Integration via Classical Morphological Features.** Shared UMAP embedding using classically measured neuronal features (e.g., branch points, tortuosity) instead of CAJAL embeddings. The overall matching accuracy decreases to 0.25.

#### S4.8 Additional results on patch-seq: Baseline comparison via supervised PCA

To contextualize GeoAdvAE's performance, we implemented a baseline comparison using a supervised linear dimensionality reduction approach. This analysis aims to determine the extent to which broad biological prior information (e.g., Excitatory vs. Inhibitory status) contributes to cross-modal alignment accuracy compared to the non-linear, geometry-aware integration of our proposed framework.

*1D Supervised PCA procedure.* Following the framework of Supervised Principal Component Analysis (SPCA) [1], we sought to identify 1D axes in both modalities that maximize dependence on the Excitatory-Inhibitory (E-v-I) spectrum. The procedure was as follows:

1. **Initial Feature Extraction:** We performed 50-dimensional PCA on the normalized gene expression (GEX) and morphological (CAJAL) data separately to reduce initial noise and stabilize the mapping.
2. **Supervised Axis Estimation:** We applied SPCA to each PCA space, using the E-v-I labels as the supervising response variable.
3. **Dimensionality Constraint:** We restricted the output to a single dimension ($d = 1$) for both modalities. This 1D projection allows for a direct, intuitive alignment along the primary axis of variation defined by the biological prior.
4. **Evaluation:** We assessed the alignment using ground-truth patch-seq pairings and computed a 1-KNN cross-modal matching metric to determine if cell types were correctly identified across modalities.

*Interpretation of baseline results.* The supervised PCA baseline, visualized in Fig. S13, achieved a cell-type matching accuracy of 28.76%. This simple linear method reaches nearly 29% accuracy, underscoring that the prior information (E-v-I status) by itself provides a critical signal for diagonal integration; this corroborates our findings in Appendix S4.1. Indeed, many complex methods in our benchmarking that lacked this prior performed significantly worse than this linear baseline.

However, our full GeoAdvAE framework achieves 34% accuracy, which is significantly higher than the 28.76% achieved by the supervised PCA. This improvement demonstrates the clear benefit of using non-linear methods and geometric regularization to resolve more granular cell-type structures. Furthermore, while SPCA is limited to a simple 1D alignment for visualization, GeoAdvAE enables a high-dimensional integrated latent space that supports advanced diagnostics, such as Monte Carlo transport mapping and integrated gradient attributions, as described in previous sections.

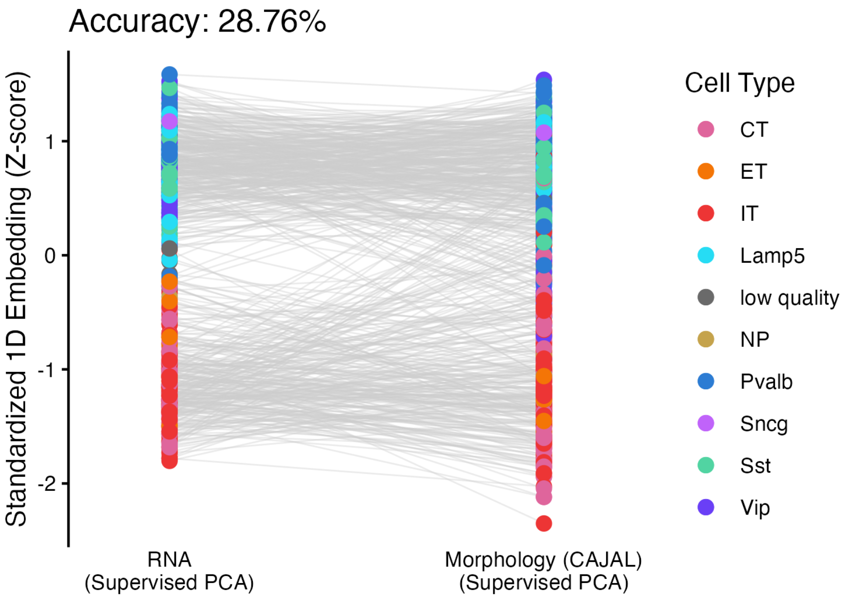


**Fig. S13. Supervised PCA Baseline.** 1D alignment of patch-seq neurons using Supervised PCA anchored by Excitatory/Inhibitory labels. Gray lines indicate ground-truth pairings. The 28.76% 1-KNN accuracy serves as a supervised linear baseline for our non-linear integration.

#### S4.9 Additional results on patch-seq: Integration performance in imbalanced modality settings

In real-world applications, it is common to encounter a significant imbalance between the number of available transcriptomic profiles and the number of reconstructed morphologies. This phenomenon is demonstrated in our microglial analysis. To evaluate the robustness of GeoAdvAE under such conditions, we utilized the full patch-seq dataset comprising 1329 neurons. While only 645 of these cells have joint gene expression (GEX) and morphological measurements (analyzed in the main text), the remaining cells possess only GEX data. This experiment aims to understand how the inclusion of additional, unpaired GEX samples from the same biological distribution affects the shared latent space and cross-modal alignment accuracy.

We applied GeoAdvAE to the imbalanced dataset (1329 GEX samples vs. 645 Morphologies), using the same hyperparameters as in the balanced 645-cell analysis. We evaluated the integration using the 1-KNN cross-modal cell-type matching accuracy, calculating both the overall mean accuracy and the modality-specific directions ($GEX \to Morph$ and $Morph \to GEX$).

The integrated latent space for the full dataset is visualized in Fig. S14. The overall 1-KNN accuracy was **27.90%**, a slight decrease from the balanced setting. However, qualitative and quantitative analyses reveal several important nuances:

- **Improved Latent Separation:** Visually, the inclusion of more GEX samples leads to a more distinct and accurate separation between excitatory (E) and inhibitory (I) neuronal families in the shared embedding.

- **Preserved GEX-to-Morph Alignment:** The $GEX \to Morph$ alignment remains highly robust at 31%. This suggests that the additional GEX information allows the model to better resolve cell states that align with known morphological phenotypes, even as the total number of cells doubles.
- **Asymmetric Accuracy:** In contrast, the $Morph \to GEX$ alignment accuracy dropped to 24.65%. As seen in Fig. S14, certain morphological samples do not cleanly map to GEX clusters, likely due to the "crowding" of the latent space by additional transcriptomic profiles that lack true morphological counterparts.

We conclude that increasing the GEX sample size improves the model's ability to prioritize biological information needed to separate states aligned with morphology. However, researchers should be aware that extreme modality imbalance can make mapping from the less-sampled modality (morphology) more challenging.

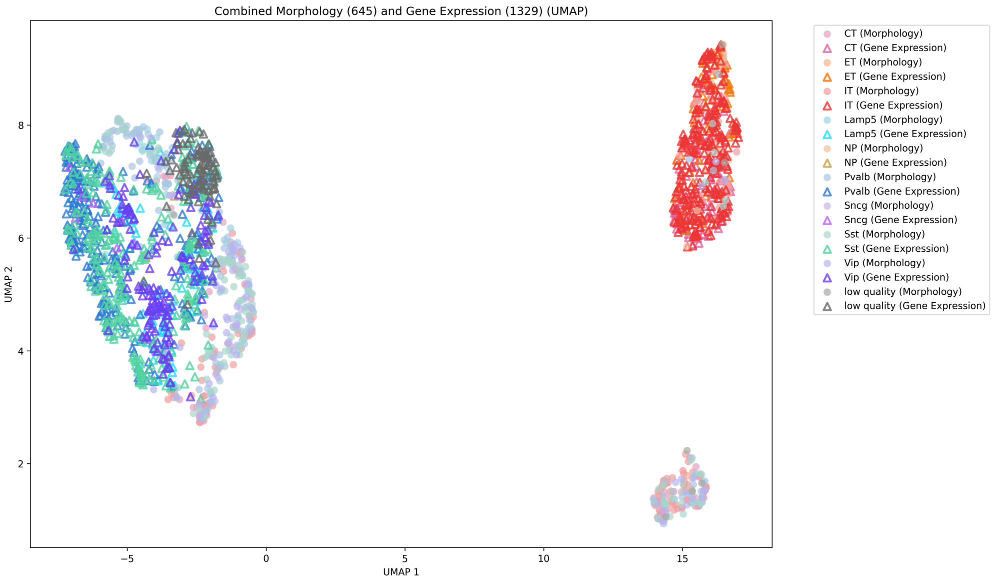


**Fig. S14. Imbalanced Modality Integration.** Shared UMAP embedding using all 1329 patch-seq GEX profiles integrated with 645 morphological reconstructions. Triangles denote GEX and circles denote morphology. The integration maintains strong modal separation and robust $GEX \to Morph$ alignment despite the sample imbalance.

##### S4.10 Additional results on patch-seq: Assessing the best-case shared structure via Tilted-CCA

To provide a benchmark for the maximum degree of shared information that could be recovered in our integration, we performed an analysis using Tilted Canonical Correlation Analysis (Tilted-CCA) [12]. Unlike diagonal integration methods like GeoAdvAE, which operate on unpaired data, Tilted-CCA is a paired method that quantifies the "intersection of information" between modalities.

Existing unsupervised integration methods typically aim to capture the "union of information" by aggregating axes of variation across modalities to improve cell-type discovery. However, as shown in Fig. 1 of the main text, the RNA modality resolves significantly more granular cell-type differences than morphology alone. Consequently, our goal is not to combine all information, but rather to isolate the specific axes of variation that are strictly shared between the two modalities. This analysis allows us to assess the theoretical "ceiling" of diagonal integration by identifying the best-case structure that an unpaired method could hope to learn.

We applied Tilted-CCA to patch-seq neurons for which ground-truth pairings are available. The procedure was implemented as follows:

1. **Input Dimensionality:** We utilized the leading 50 principal components (PCs) of the gene expression matrix and the leading 50 PCs derived from the CAJAL Gromov-Wasserstein distance matrix.
2. **Common Score Computation:** We decomposed the canonical score matrices to compute a 50-dimensional common score matrix. This process involved constructing a nearest-neighbor graph for each modality (using $k = 20$ neighbors) and iteratively fine-tuning the "tilt" of each latent dimension to approximate the intersection of the modality-specific manifolds.
3. **Visualization:** The resulting 50-dimensional common scores, which represent information present in both RNA and morphology by definition, were projected into 2D using UMAP for visualization.

The Tilted-CCA common space embedding is visualized in Fig. S15.

– **High-Level Separation:** We observe a clear and robust separation between excitatory and inhibitory neuronal families, maintaining the color scheme established in the main text.
– **Morphological Separability:** More importantly, this shared embedding provides strong evidence that morphological features are indeed separable by cell type when isolated via their coordination with transcriptomic signals.

Because this structure is mathematically restricted to the intersection of the two data matrices, it defines the limit of what a diagonal integration method can realistically achieve. The strong cell-type clusters observed here validate the presence of sufficient shared geometric structure between the intricate forms of neural cells and their molecular states to justify cross-modal alignment.

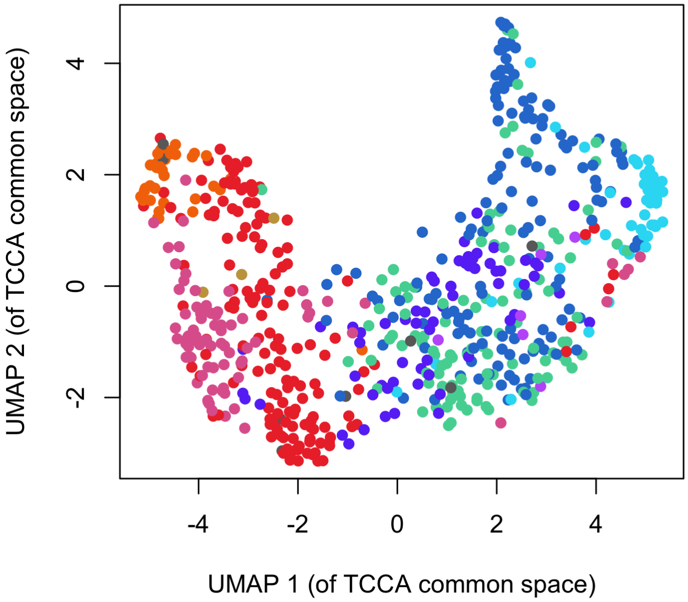


**Fig. S15. Best-Case Shared Latent Structure via Tilted-CCA.** UMAP visualization of the 50-dimensional common score matrix estimated from paired patch-seq data. This embedding represents the theoretical ceiling for diagonal integration, demonstrating that cell-type differences are encoded in the shared variation of RNA and morphology. The coloring of cells is based on the coloring scheme in Fig. 1 in the main text.
